# Out of sight, out of mind: Removal of predators by fisheries alters reef fish behaviour

**DOI:** 10.64898/2025.12.08.692929

**Authors:** Shawn Dsouza, Akshta Joshi, Maria Thaker, Kartik Shanker

## Abstract

Pervasive human activities have led to the extirpation of predator species in many ecosystems. Understanding the extent and mechanisms by which such predator loss influences prey behaviour is critical for predicting the ecological consequences of trophic downgrading. We investigated reef fish behaviour in response to predator presence within and outside a protected area in the South Andaman Islands, India. We used 3D printed models to simulate the presence of two different types of predators; a sit-and-wait predator, a grouper (*Mycteroperca rosacea*) and a wide-ranging pursuit predator, a barracuda (*Sphyraena barracuda*). We recorded foraging, vigilance, and movement behaviours of herbivorous and invertivorous fish encountered in each plot. Overall, we found that fish were significantly more vigilant in response to both predator models compared to a non-predator control within protected areas but not outside protected areas. Foraging behaviour was suppressed by the predator treatments; however, herbivorous and invertivorous fish responded differently within protected areas. Fish movement behaviours also differed across predator treatments, prey foraging guilds, and areas. Anti-predator responses are thus highly contingent on habitat and both predator and prey traits. We highlight how the selective removal of predators by humans alters predation risk for prey and hence their risk-sensitive behaviours.

## Introduction

Predators are known to play an important role in ecosystems through their top-down control of prey populations and behaviour [1]. In addition to their ecological value, predator species also hold significant economically valuable and are, thus, disproportionately targeted by hunters and fishers [2]. This pressure has driven the decline or extirpation of top predators from many parts of the globe [3]. This loss of predators has had large scale impacts on the ecology and economies of many regions (e.g. [4–6]). The direct consequences of predator loss, such as mesopredator release, are well documented [7,8]. The removal of predators can also disrupt many ecological interactions, leading to cascading effects that are wide ranging and include changes to ecosystem function – a process known as trophic downgrading (sensu [1]).

The behavioural effects of the loss of predators on prey have been hypothesised to be just as important as the density dependent effects on them [9]. Removal of predators by human hunters and fishers can lead to reduced risk for prey in these ecosystems and may thus alter prey behaviour [10–12]. Over time, reduced risk may lead to a loss of predator recognition and antipredator behaviour [e.g. 20]. The effects of altered prey behaviour may also cascade to lower trophic levels, altering plant communities and affecting ecosystem processes [14,15]. For example, changes in herbivore foraging behaviour due to reduced predator abundance have been shown to alter seagrass distribution in coral reefs [16,17].

Thus, it is important to consider how prey perceive and respond to the risk of predation in their environment [18–20]. Prey are known to induce a variety of developmental, morphological, physiological, and behavioural responses to reduce the risk of depredation [20,21]. Some of these responses can have energetic or fitness costs. Behaviourally, prey can implement strategies such as increased vigilance, which increases their probability of detecting predators, often at the cost of foraging, or they can reduce movement to avoid being detected by predators [22]. Prey may also choose to forage in less risky patches at the cost of resource quality in order to avoid predators altogether [23,24]. The utilisation and allocation of behavioural responses depend on the nature of predation risk, how intense and variable it is, and the durations that prey are exposed to it. For example, prey exposed to short-term but intense predation risk may choose to not forage for that duration; however, when predation risk is chronic, the internal states of prey such as hunger or mating status may determine their behaviour [25]. Prey traits, such as foraging mode and food choice, may also influence their exposure to predation risk. For example, prey species that actively search for food across large areas, such as insectivores, often experience higher exposure to predators as they cover broader habitats compared to more sedentary prey like herbivores [26].

In addition to prey traits, the response of prey to predation risk may be determined by the characteristics of the habitat they occupy and the traits of the dominant predators in those habitats [27]. The threat of different types of predators may also vary by habitat. For example, sit-and-wait predators like groupers (Epinephelidae) rely on highly structured habitats for cover in order to stealthily ambush their prey. On the other hand, wide-ranging pursuit predators like barracuda (Sphyraenidae) often prefer open habitats where they can more easily spot and chase prey [28]. The behavioural response of prey to these distinct types of predators depends on the habitat and their abundance in that environment [13]. Consequently, the selective removal of different types of predators from a given system may have divergent effects of the behaviour of prey species and individuals.

Here, we aim to investigate the behavioural effects of the loss of piscivorous predators on herbivorous and invertivorous fish in a nearshore coral reef ecosystem in the Andaman Islands of India. We hypothesised that fish outside protected areas, where predator populations are reduced due to fishing pressure, will exhibit a diminished response to predator models, suggesting a potential loss of anti-predator behaviour due to reduced exposure to natural predation risk. To this end, we conducted a predator cue experiment wherein we exposed reef-associated fish to two different model predators positioned on the reef floor (Figure 1). The predator models represent typical reef predators with ambush (grouper) and coursing (barracuda) hunting strategies, allowing us to determine prey responses to different predator types [28,29], Figure 1). We recorded the behaviours of fish that entered the field of view in each sampling plot, and quantified their foraging, movement, and vigilance responses. We also included algal cover, substrate rugosity, and the grouping status of prey in our analysis as previous studies have shown their influence on fish behaviour [30]. We compared these behaviours across treatments within and outside protected areas to determine differences in responses across predator identity and protection status.

**Figure 1:**
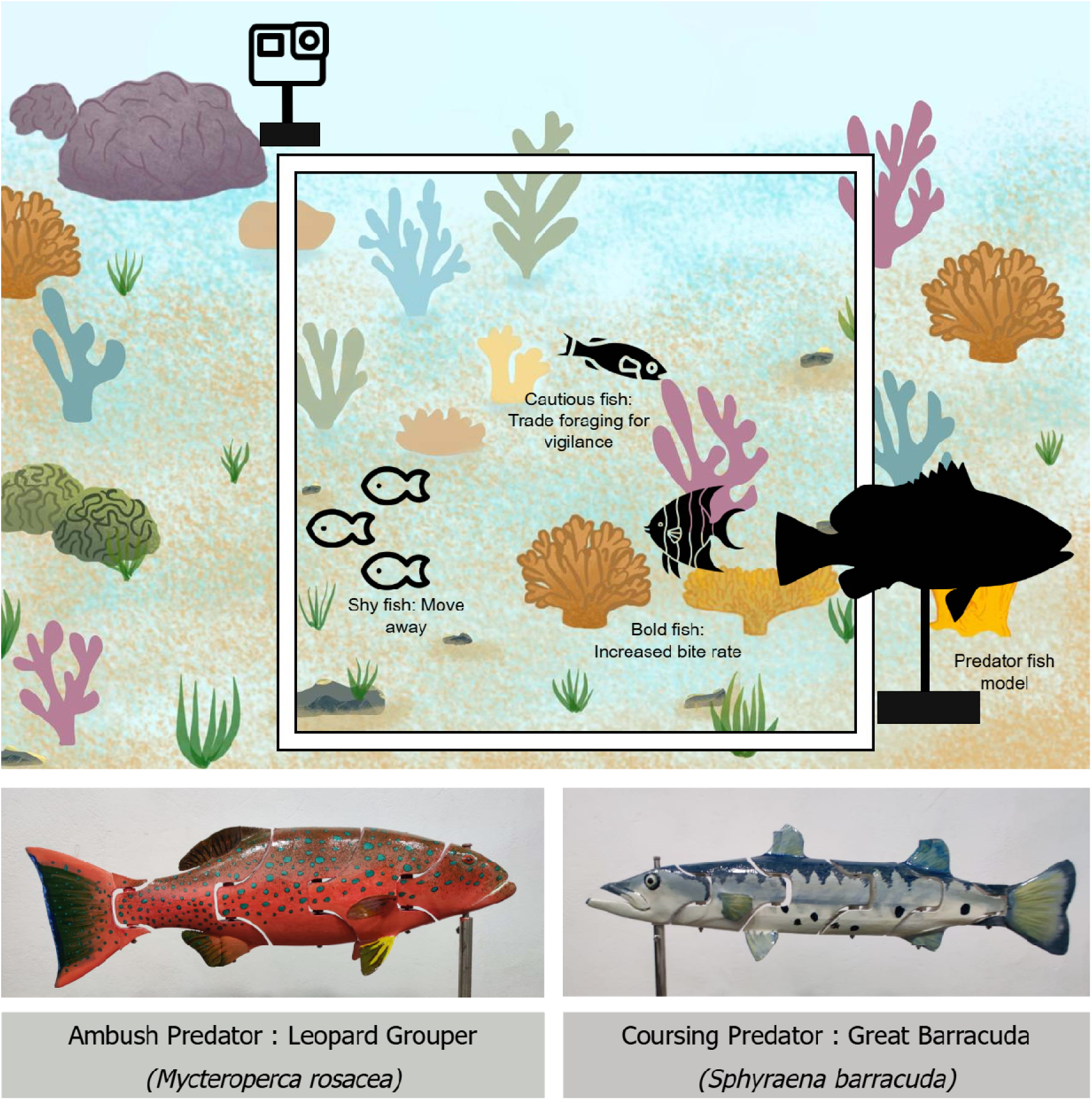
Experimental setup for predator-cue trials. A camera was placed on a 1×1 m plot, with a treatment, positive control, or negative control positioned in the opposite corner. Each plot was filmed for 1 hour. Four plots were deployed per day, each randomly assigned to one of four treatments: ambush predator, coursing predator, positive control, or negative control. The experiment was repeated five times both inside and outside the Mahatma Gandhi Marine National Park (MGMNP). *Bottom:* 3D-printed predator models used in the in-situ experiments.

## Methods

### Site selection

We conducted our experiments at Boat Island within Mahatma Gandhi Marine National Park (MGMNP) and in Allan’s patch adjoining the protected area. Fisheries in the Andaman and Nicobar Islands (ANI) primarily target predatory fish such as groupers. A previous study has shown that predator abundances within MPAs in the region were on average 1.5 times greater than outside MPAs [30]. Allan’s patch experiences moderate fishing pressure, predominantly through gillnets and hook-and-line methods [31]. MGMNP enforces no-take fishing restrictions within the park boundaries, making this area an ideal control setting for testing our hypotheses.

### In-situ experiments

We deployed four treatments both inside and outside the protected area (n = 5 each): a negative control, a positive control, a leopard grouper (*Mycteroperca rosacea*) model, and a great barracuda (*Sphyraena barracuda*) model. All four treatments were deployed simultaneously on a given day by randomly placing four 1×1Lm plots over reef patches that varied in rugosity and resource availability. Treatments were randomly assigned to plots, and plots were placed a minimum of 5m apart. A GoPro camera was mounted at one corner of each plot, providing a field of view of approximately 2Lm [28,32]. In the negative control treatment, only the camera was placed on the plot. In the positive control, a hollow tube approximately the same length as the predator models was placed opposite the camera. For the predator treatments, the corresponding model was positioned on the corner of the plot diagonally opposite the camera. Each camera recorded continuously for 1 hour, resulting in a total of 20 hours of video recordings per fishing pressure level. All behavioural observations were conducted between 8:00 AM and 12:00 PM to account for potential changes in fish activity through the day [33].

### Measuring plot-level co-variates: resource availability and habitat complexity

We took a top-down photograph of each plot with the quadrat placed on the reef bed, and used Coralnet [34] to determine the relative cover (%) of corals, sponges, algae and other substrates (sand, rock, rubble). Following the experiment, we measured rugosity at three different angles around each camera plot using the chain transect method [34]. Rugosity is the estimated ratio of length covered on the ground to the total length of the chain (190 m) [35].

### Video processing for fish behavioural observations

Fish responses to our treatments were scored from the videos recorded from each plot. We discarded the first 10 minutes of each recording to account for any disturbance caused during the deployment of the cameras. We then processed five random 2-minute samples of the remaining video for a total of 10 minutes sampled at each plot. We identified and counted the number of all fish species within those recordings. We visually estimated the size class of sampled individuals (within 10 cm) using the models as reference [28,29].

We quantified the behavioural traits for each focal individual that entered our plot during the observation window. Behaviours were classified into foraging, vigilance, and movement (Table 1). We calculated individual time budgets as the proportion of total observation time spent in each behavioural state. This was estimated for individuals observed for more than 45s. When individuals were observed foraging, we additionally counted all bites within each foraging bout to estimate bite rates (number of bites per second).

**Table 1:**
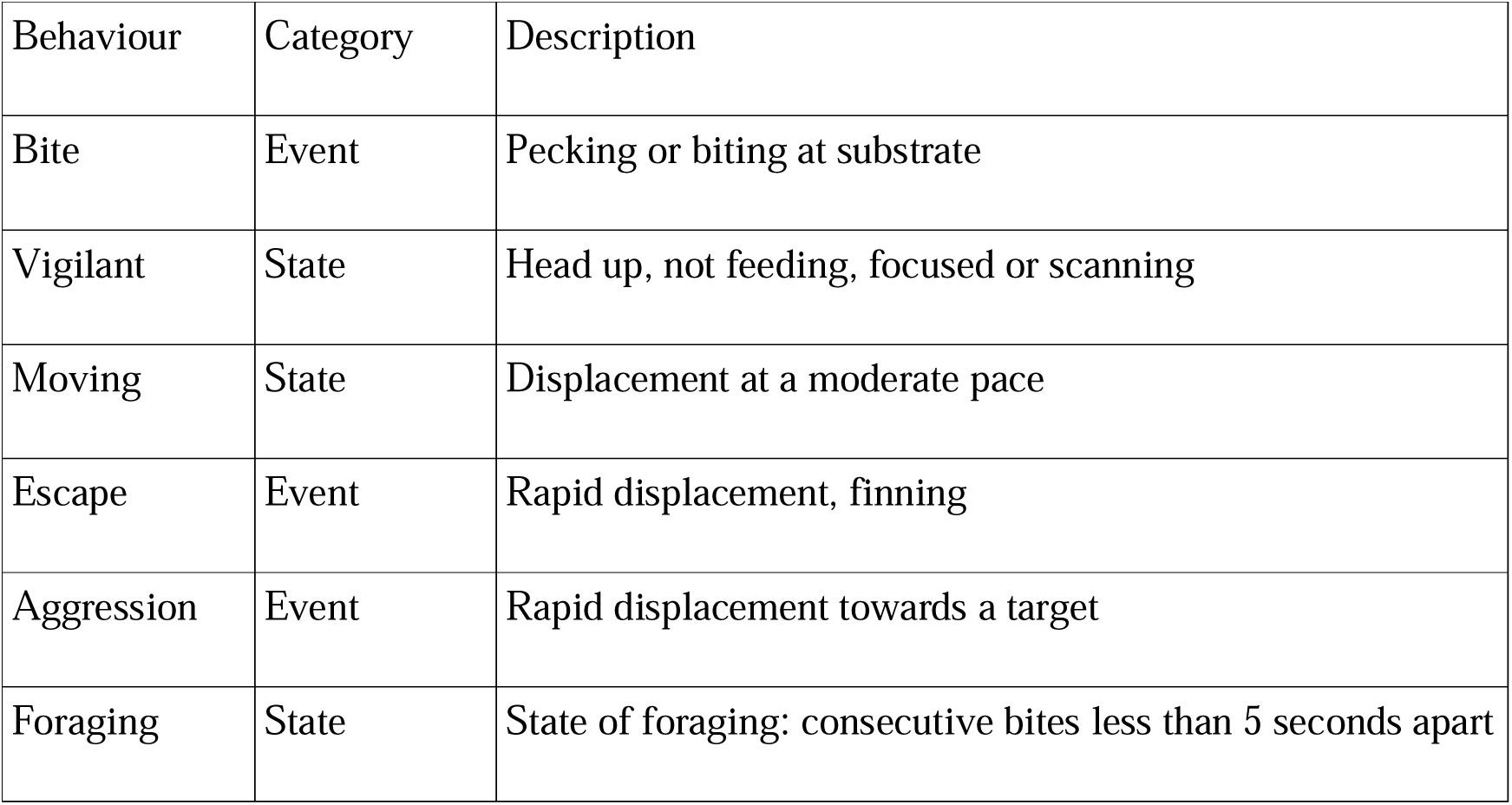
Ethogram describing definitions of behaviours utilised for behavioural assays.

| Behaviour | Category | Description |
| --- | --- | --- |
| Bite | Event | Pecking or biting at substrate |
| Vigilant | State | Head up, not feeding, focused or scanning |
| Moving | State | Displacement at a moderate pace |
| Escape | Event | Rapid displacement, finning |
| Aggression | Event | Rapid displacement towards a target |
| Foraging | State | State of foraging: consecutive bites less than 5 seconds apart |

We also noted any aggressive behaviour by the focal individuals against natural predators, conspecifics, and heterospecifics. We did not observe any predation events or any disturbance due to human activities such as fishing or SCUBA diving. Additionally, we noted whether individuals were foraging in a group, and recorded group composition.

### Analysis

We used a causal inference framework along with Bayesian estimation to model the response of herbivorous and invertivorous fish individuals to predator models [36,37]. We used counter-factual simulations based on fitted models to estimate the effect of predictors by simulating outcomes under unobserved conditions (McElreath, 2018). We used weakly informative, sceptical priors for all intercepts and coefficients (*Normal(0, 0.5)*), and appropriate weak priors for standard deviations, scale, precision, and shape parameters (*Exponential(2)*). For each estimate, we report the posterior probability of superiority—the probability that a parameter is greater than zero. Values near 0.5 indicate little or no effect, while values above 0.9 or below 0.1 indicate strong positive or negative effects, respectively. We assessed model fit using Bayesian R^2^ [38]. All analyses were conducted in R 4.5.0 using the brms package [39,40].

Proportion of time spent on each behaviour was modelled using an ordered beta regression to account for zero- and one-inflation [41]. Predictors included treatment (negative control, positive control, barracuda or grouper), protection (inside or outside MPA), algal cover, rugosity, grouping status (solitary, grouping), foraging guild (piscivore, herbivore and invertivore), and size class (0-10cm, 10-20cm, … >80cm). We included full interactions between treatment, protection and foraging guild. The day of the deployment and species (nested within family) were included as random effects. We calculated the effect of barracuda and grouper treatments relative to control as log odds ratios (LOR). For individuals that foraged, bite rates were modelled as a gamma process with a log link. Predictors and random effects were as above, and effects were calculated as log response ratios (LRR).

We found that fish did not change the proportion of time foraging and moving in response to the positive control relative to negative control. However, fish were overall significantly less vigilant, herbivores decreased foraging rate, and invertivores increased foraging rate in the positive control plots compared to the negative control plots (Table 1, 2 of Appendix D). We thus used the positive control as the contrast for the barracuda and grouper model treatments.

## Results

We collected a total of 40 plot-hours of video across the four treatments within and outside MGMNP. We encountered 431 individuals belonging to 105 fish species during the behavioural assays. Of these, 70 species were invertivores (n = 336), 39 species were herbivores (n = 202), 17 species were piscivores (n = 63) and 21 species (n = 170) were assigned to more than one guild. Piscivores were excluded from our analysis of behaviour. We observed 126 individuals as part of groups. Additionally, we observed 18 instances of predator avoidance behaviour in response to model predators and 16 instances of conspecific aggression behaviour.

### Effect of predator models on fish foraging behaviour

We found that herbivorous fish did not significantly alter their foraging time (*P_inside_* =0.381, *P_outside_* =0.416) or their foraging rates (*P_inside_* =0.323, *P_outside_* =0.357) in response to the barracuda model relative to positive control model, both within and outside protected areas (Table 3 & 4 of Appendix D, Figure 2). However, herbivores reduced the proportion of time they spent foraging in response to the grouper model (*P_inside_* =0.232, *P_outside_* =0.175, Figure 2). Herbivores also increased their foraging rates in response to the grouper model within the MPA but not outside the MPA (*P_inside_* =0.834, *P_outside_* =0.599; Figure 2).

**Figure 2:**
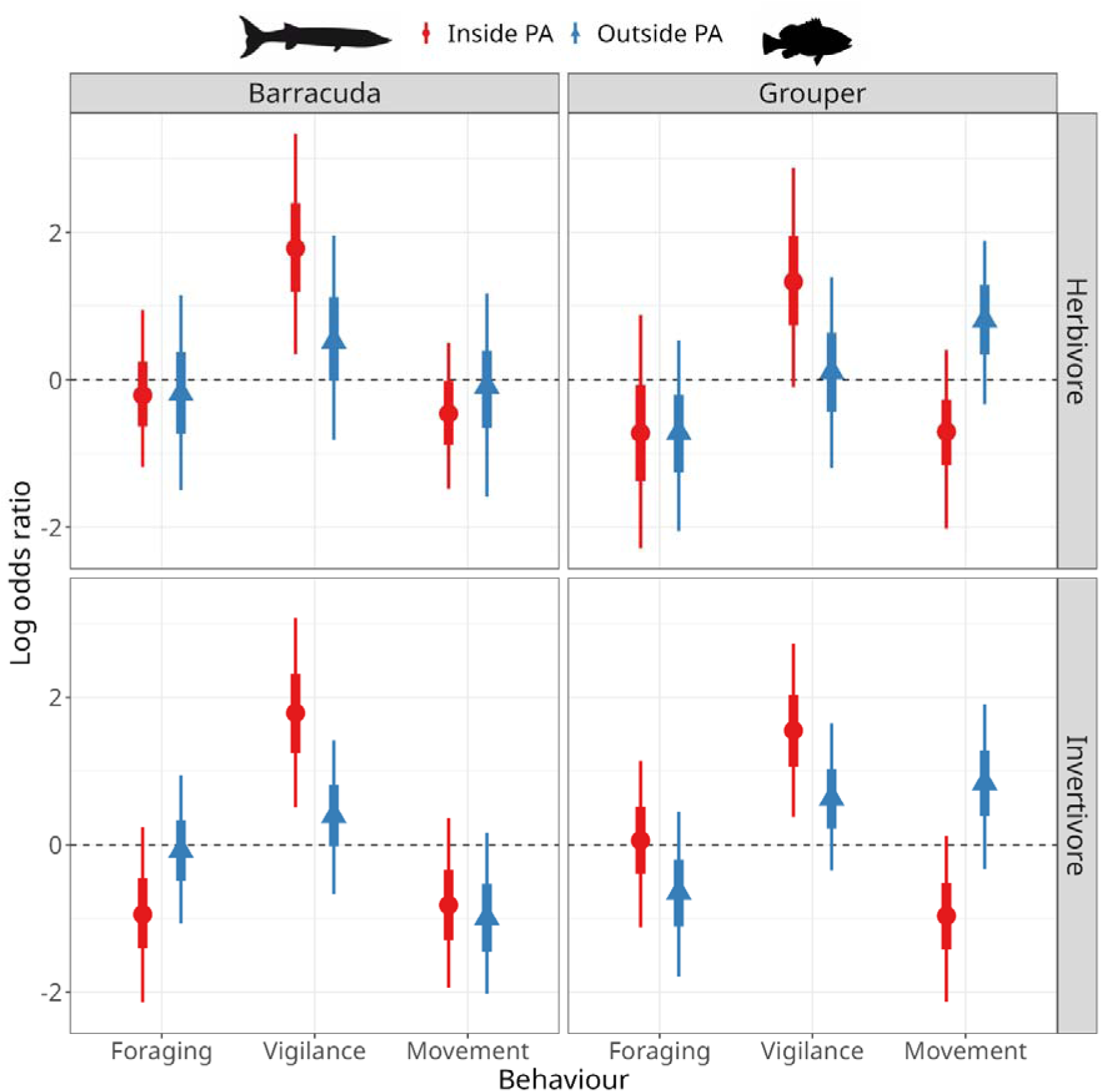
Effect (log odds ratio) of predator model treatments on the proportion of time herbivorous and invertivorous fish spent foraging, moving, and vigilant within (red circles) and outside (blue triangles) protected areas. Values above zero represent higher response relative to the positive control and values below zero represent a lower response.

Invertivores significantly reduced foraging time in response to barracuda models inside the MPA but not outside the MPA (*P_inside_* =0.095, *P_outside_* =0.454). Additionally, invertivores significantly reduced their foraging rates in response to barracuda models both within and outside protected areas (Figure 3). This effect was more pronounced within protected areas compared to outside (*P_inside_* =0.008, *P_outside_* =0.055). Invertivores foraged for shorter times in response to grouper models outside the MPA but not inside the MPA (*P_inside_* =0.538, *P_outside_* =0.162). However, invertivores did not alter their foraging rate in response to grouper models (*P_inside_* =0.376, *P_outside_* =0.312, Figure 3).

**Figure 3:**
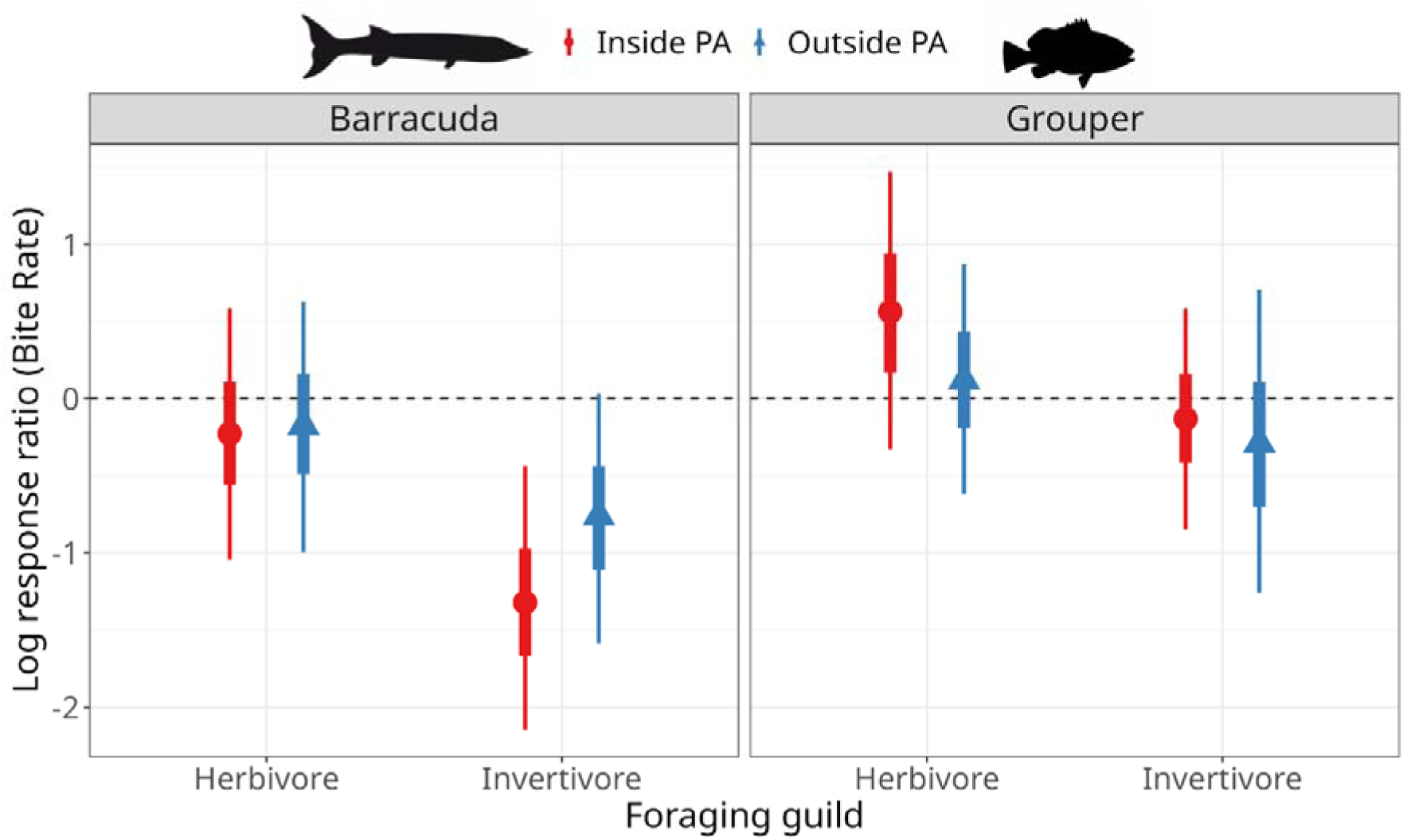
Effect (log response ratio) of predator model treatments relative to positive control on the bite rates (per second) of herbivorous and invertivorous fish within and outside protected areas.

### Effect of predator models on fish vigilance behaviour

Herbivores were significantly more vigilant in barracuda treatment plots compared to control plots. The effect was more pronounced within the MPA compared to outside the MPA (*P_inside_* =0.980, *P_outside_* =0.739). Similarly, herbivores spent more time being vigilant in grouper treatment plots within the MPA but not outside the MPA (*P_inside_* =0.934, *P_outside_* =0.561). Invertivores were also significantly more vigilant in barracuda treatment plots within the MPA but not outside the MPA (*P_inside_* =0.989, *P_outside_* =0.739). Additionally, invertivores were significantly more vigilant in grouper treatment plots. The effect was once again more pronounced within MPA compared to outside the MPA (*P_inside_* =0.985, *P_outside_* =0.851, Figure 2).

### Effect of predator models on fish movement behaviour

The proportion of time herbivores spent moving did not change in response to the barracuda model (*P_inside_* =0.240, *P_outside_* =0.448). However, invertivores moved for significantly less time in response to barracuda models both within and outside the MPA (*P_inside_* =0.076, *P_outside_* =0.890). In response to the grouper model, both herbivores and invertivores significantly reduced time spent moving within the MPA. Conversely, both herbivores and invertivores increased movement in response to grouper models outside the MPA (Table 3 of Appendix D, Figure 2).

### Effect of habitat complexity, resource availability, and grouping on fish behaviour

Herbivorous and invertivorous fish spent significantly less time foraging as plot rugosity increased (*β = -0.067 [ -0.388,0.331], P(β > 0) = 0.025*), but fish foraged at significantly higher rates as rugosity increased (*β = -0.169 [-0.398, 0.052], P(β > 0) = 0.10*). Fish were more vigilant (*β = 0.080 [-0148, 0.315], P(β > 0) = 0.720*) and spent more time moving in areas with higher rugosity (*β = 0.156, [-0.074,0.374], P(β > 0) = 0.879*). Biomass cover did not influence the duration of foraging (*β = -0.068 [-0.314,0.172], P(β > 0) = 0.325*), but fish tended to forage at a slightly higher rate when biomass cover was greater (*β = -0.113 [-0.309, 0.09], P(β > 0) = 0.174*). Fish spent less time moving in areas with higher biomass cover (*β = -0.053 [-0.286, 0.168], P(β > 0) = 0.339*), but time spent vigilant was unaffected (*β = - 0.005 [-0.230, 0.220], P(β > 0) = 0.485*). When in groups, fish spent significantly less time vigilant (*β = -0.205 [-0.811,0.392], P(β > 0) = 0.286*) and moved more (*β = 0.252 [-0.066, 0.563], P(β > 0) = 0.913*). However, grouping did not affect foraging duration (*β = -0.328 [-0.589,-0.052], P(β > 0) = 0.343*) or foraging rate (*β = -0.029 [-0.411, 0.352], P(β > 0) = 0.453*).

## Discussion

In the present study, we aimed to investigate whether the removal of piscivorous predators by fisheries in the Andaman Islands has had measurable impacts on the behaviour of reef-associated fishes. Specifically, we expected that fish outside marine protected areas (MPAs), where predator abundances are lower due to sustained fishing pressure, would show weaker behavioural responses to predator models than fish inside MPAs. Most notably, fish within the MPA were significantly more vigilant in response to both types of predator models relative to controls; however, fish outside the MPA did not alter their vigilance behaviour in response to the predator models. Fish foraging behaviour was suppressed by both the barracuda and grouper models, but the degree of change varied by prey foraging guild and protection status of the habitat (Figure 3). Similarly, fish also altered their movement in response to these predator models but with different directions and magnitude depending on their foraging guild and whether they were inside or outside the protected area (Figure 2).

### Effect of predator loss on prey behavioural responses

Many prey species implement behavioural anti-predator strategies to mitigate the risk of depredation [42,43]. However, the removal of predators by human hunters and fishers can reduce the local intensity of predation risk experienced by prey individuals. Thus, prey in areas with higher anthropogenic pressure on predators may be able to allocate their time to different behaviours than prey individuals in areas with low anthropogenic fishing pressure. We found that both herbivore and invertivore prey responded to both predator models by increasing vigilance when they were inside the MPA, but they did not respond to the predator models outside the park. Thus, MPAs may preserve the ecological interactions that underpin risk perception in prey. Robust predator populations inside MPAs seem to maintain the selective pressure for anti-predator responses [44]. The weakened responses observed outside MPAs may be explained by prey naivety to predators, where prey lose the ability to recognize or appropriately respond to predators due to reduced exposure [13,45].

Our results are consistent with the predation risk allocation hypothesis, predicting that anti-predator behaviours are not uniformly expressed, but are shaped by the type and frequency of predator-prey encounters [20,25]. In predator-rich environments, prey devote time to vigilance, incurring a missed opportunity cost on foraging as an adaptive trade-off [23]. The absence of such trade-offs outside MPAs suggests that prey may prioritise foraging over risk avoidance, potentially enhancing individual growth or reproduction in the short term; but this response may potentially leave populations poorly adapted to cope with predation risk if predator abundances recover faster than antipredator responses regain [46]. Similar shifts in prey behaviour have been reported in other reef studies, where predator depletion through overfishing alters grazing intensity and habitat use, sometimes triggering cascading effects on reef benthic communities [47,48]. For instance, altered herbivory patterns due to a lack of predator-induced fear can shift the competitive balance between corals and algae [49]. Our findings therefore provide empirical support for the idea that behavioural pathways are a key mechanism through which predator extirpation reshapes ecosystem function.

### Effect of predator and prey traits on behaviour responses

One of the most striking outcomes of our experiment was the different responses of reef-associated fish to ambush and pursuit predators. Herbivores demonstrated stronger responses to the grouper model compared to the barracuda model, significantly reducing foraging time and increasing their foraging rates (Figure 2 & 3). Invertivores, on the other hand, exhibited stronger behavioural adjustments to barracuda models, including significant reductions in foraging rates in both protected and fished reefs (Figure 3). Predator impacts may thus be contingent on prey traits, with prey individuals perceiving different levels of risk depending on their foraging strategy and food preference [27]. Herbivores typically forage in complex habitats with high rugosity, making them more vulnerable to ambush predators like groupers. Thus, grouper cues may be perceived as more ecologically relevant or threatening to herbivores. By contrast, invertivores are known to forage in open sandy patches making them more vulnerable to pursuit predators like barracuda [27]. Such predator-specific responses highlight the importance of functional diversity of piscivores in coral reef ecosystems in maintaining complex trophic interactions [49–51].

### Effect of habitat characteristics and grouping on anti-predator responses

As expected, habitat characteristics can alter prey responses to predators. For example, refuge availability can reduce predation risk for prey, thereby changing the spatial distribution of risk at the local scale [52]. As reef structure became more complex, we found that fish reduced the time spent foraging but increased the rate at which they forage. More complex habitats also elicited greater vigilance and movement, suggesting that prey perceived higher risk in structurally complex areas. Prey internal state such as hunger may also alter the amount of time they allocate to different behaviours. Fish did not change the proportion of time spent on foraging or vigilance with increasing biomass cover, but they did forage at higher rates. This suggests that the trade-off between foraging and vigilance in time was maintained, while foraging effort was increased.

Prey behaviours such as grouping also influence the overall risk experienced by prey individuals [53]. In our plots, fish in groups spent less time being vigilant and moved more frequently than solitary individuals. Solitary fish, in contrast, foraged less and allocated more time to vigilance, consistent with the dilution effect and advantages of collective defence observed across taxa [54]. These patterns indicate that grouping functions as a behavioural buffer against predation risk, allowing individuals to allocate effort from vigilance to foraging.

## Conclusion

Our experiment revealed that prey responses to predation risk may be altered by the loss of predators due to human disturbance. In addition, prey behavioural responses are highly context dependent on predator and prey traits, such as predator hunting mode and prey foraging preferences, highlighting the role of functional diversity in predator assemblages. Within MPAs, where predator populations remain relatively intact, reef fishes display strong trade-offs between foraging and vigilance in response to predator cues, consistent with intact predator-prey dynamics. Outside MPAs, where fishing has reduced predator densities, fish responses are diminished, suggesting a loss of predator recognition and risk sensitivity. Thus, the ecological benefits of MPAs extend beyond population recovery of predators and encompass the preservation of key behavioural processes that regulate ecological interactions.

## Acknowledgements

We would like to thank all the staff at Andaman Nicobar Environment Team (ANET), especially our boat captains and crew; Saw Thesorow, Jeevan Horo, Babu Kutty, Saw Watha (Agu), Anand James Tirkey, and Sebian Horo, for their logistical support. We thank Chaitanya Arjunwadkar his assistance with field work and suggestions on methods, and Samar Ahmad and Nina Simon for their assistance during field work. Finally, we would like to thank the ANI forest department, the MGMNP ranger office, the RJMNP ranger office, and ANI Fisheries Department for providing us with permits to conduct our work.

## Data accessibility

Data and code for analysis are available at https://doi.org/10.5281/zenodo.17854841

## Appendix A: List of fish species sampled in predator model experiments

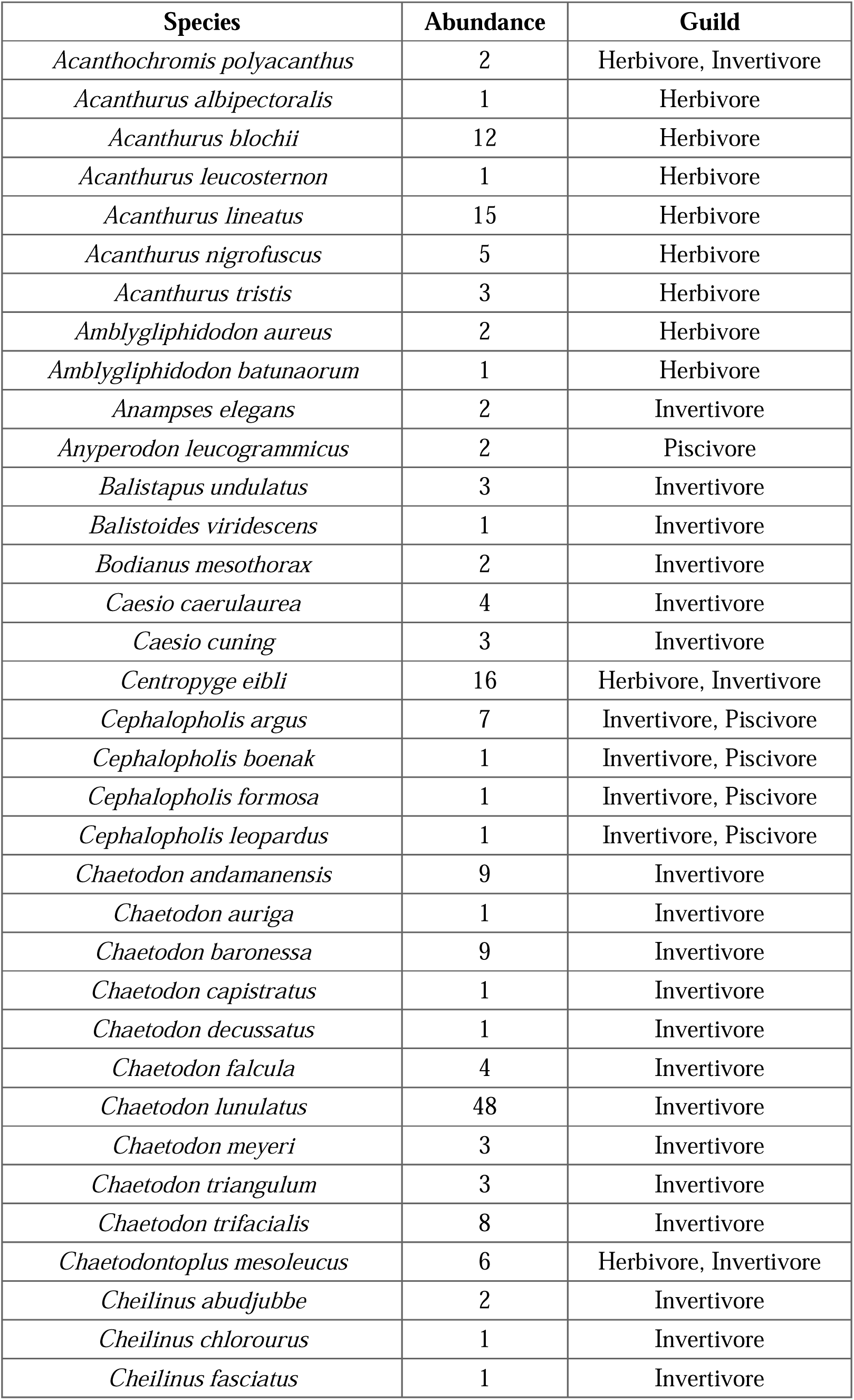
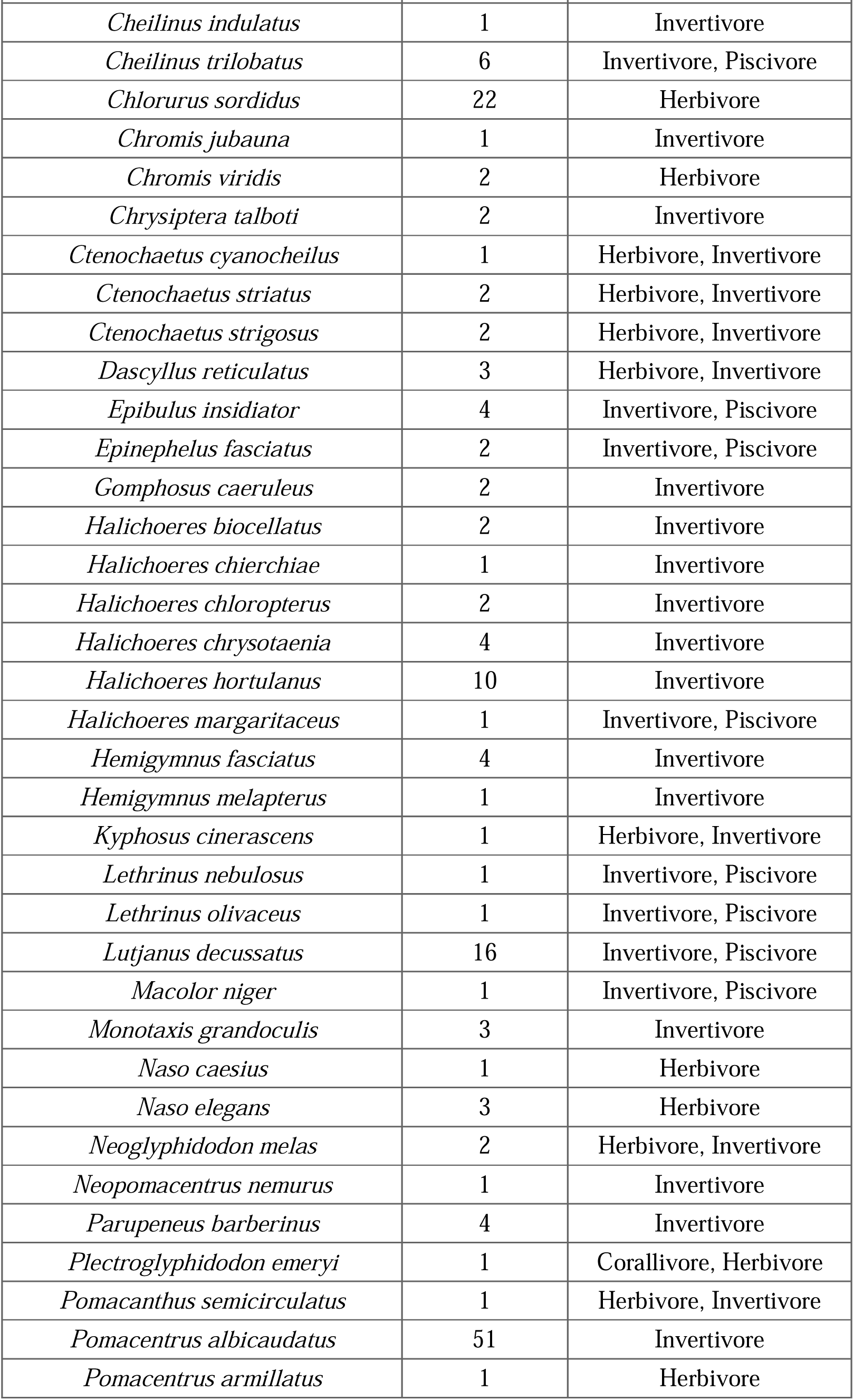
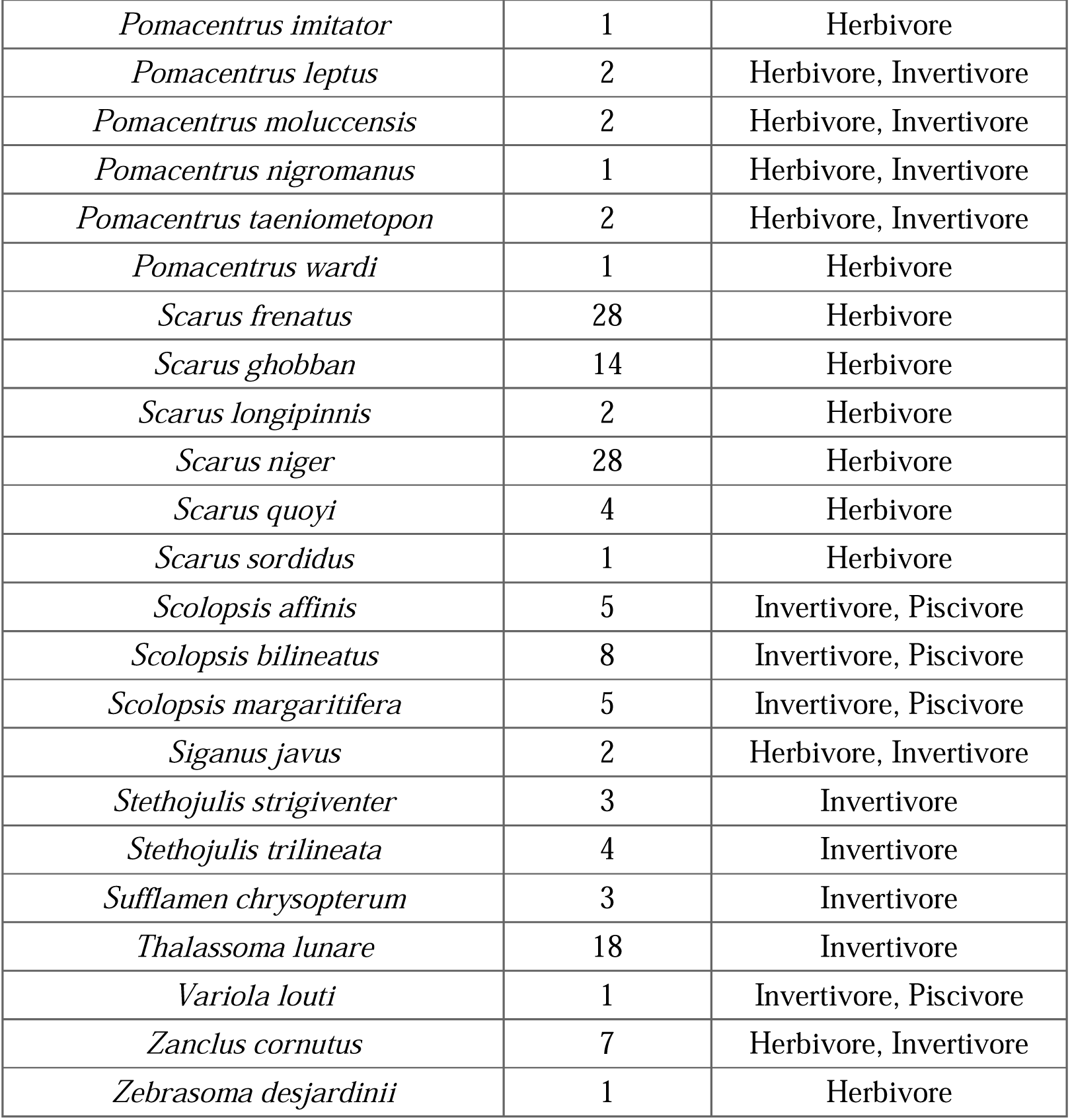

**Appendix A: List of fish species sampled in predator model experiments.**
| **Species** | **Abundance** | **Guild** |
| --- | --- | --- |
| *Acanthochromis polyacanthus* | 2 | Herbivore, Invertivore |
| *Acanthurus albipectoralis* | 1 | Herbivore |
| *Acanthurus blochii* | 12 | Herbivore |
| *Acanthurus leucosternon* | 1 | Herbivore |
| *Acanthurus lineatus* | 15 | Herbivore |
| *Acanthurus nigrofuscus* | 5 | Herbivore |
| *Acanthurus tristis* | 3 | Herbivore |
| *Amblygliphidodon aureus* | 2 | Herbivore |
| *Amblygliphidodon batunaorum* | 1 | Herbivore |
| *Anampses elegans* | 2 | Invertivore |
| *Anyperodon leucogrammicus* | 2 | Piscivore |
| *Balistapus undulatus* | 3 | Invertivore |
| *Balistoides viridescens* | 1 | Invertivore |
| *Bodianus mesothorax* | 2 | Invertivore |
| *Caesio caerulea* | 4 | Invertivore |
| *Caesio cuning* | 3 | Invertivore |
| *Centropyge eibli* | 16 | Herbivore, Invertivore |
| *Cephalopholis argus* | 7 | Invertivore, Piscivore |
| *Cephalopholis boenak* | 1 | Invertivore, Piscivore |
| *Cephalopholis formosa* | 1 | Invertivore, Piscivore |
| *Cephalopholis leopardus* | 1 | Invertivore, Piscivore |
| *Chaetodon andamanensis* | 9 | Invertivore |
| *Chaetodon auriga* | 1 | Invertivore |
| *Chaetodon baronessa* | 9 | Invertivore |
| *Chaetodon capistratus* | 1 | Invertivore |
| *Chaetodon decussatus* | 1 | Invertivore |
| *Chaetodon falcula* | 4 | Invertivore |
| *Chaetodon lunulatus* | 48 | Invertivore |
| *Chaetodon meyeri* | 3 | Invertivore |
| *Chaetodon triangulum* | 3 | Invertivore |
| *Chaetodon trifacialis* | 8 | Invertivore |
| *Chaetodontoplus mesoleucus* | 6 | Herbivore, Invertivore |
| *Cheilinus abudjubbe* | 2 | Invertivore |
| *Cheilinus chlorourus* | 1 | Invertivore |
| *Cheilinus fasciatus* | 1 | Invertivore |
| *Cheilinus indulatus* | 1 | Invertivore |
| *Cheilinus trilobatus* | 6 | Invertivore, Piscivore |
| *Chlorurus sordidus* | 22 | Herbivore |
| *Chromis jubauna* | 1 | Invertivore |
| *Chromis viridis* | 2 | Herbivore |
| *Chrysiptera talboti* | 2 | Invertivore |
| *Ctenochaetus cyanocheilus* | 1 | Herbivore, Invertivore |
| *Ctenochaetus striatus* | 2 | Herbivore, Invertivore |
| *Ctenochaetus strigosus* | 2 | Herbivore, Invertivore |
| *Dascyllus reticulatus* | 3 | Herbivore, Invertivore |
| *Epibulus insidiator* | 4 | Invertivore, Piscivore |
| *Epinephelus fasciatus* | 2 | Invertivore, Piscivore |
| *Gomphosus caeruleus* | 2 | Invertivore |
| *Halichoeres biocellatus* | 2 | Invertivore |
| *Halichoeres chierchiae* | 1 | Invertivore |
| *Halichoeres chloropterus* | 2 | Invertivore |
| *Halichoeres chrysotaenia* | 4 | Invertivore |
| *Halichoeres hortulanus* | 10 | Invertivore |
| *Halichoeres margaritaceus* | 1 | Invertivore, Piscivore |
| *Hemigymnus fasciatus* | 4 | Invertivore |
| *Hemigymnus melapterus* | 1 | Invertivore |
| *Kyphosus cinerascens* | 1 | Herbivore, Invertivore |
| *Lethrinus nebulosus* | 1 | Invertivore, Piscivore |
| *Lethrinus olivaceus* | 1 | Invertivore, Piscivore |
| *Lutjanus decussatus* | 16 | Invertivore, Piscivore |
| *Macolor niger* | 1 | Invertivore, Piscivore |
| *Monotaxis grandoculis* | 3 | Invertivore |
| *Naso caesius* | 1 | Herbivore |
| *Naso elegans* | 3 | Herbivore |
| *Neoglyphidodon melas* | 2 | Herbivore, Invertivore |
| *Neopomacentrus nemurus* | 1 | Invertivore |
| *Parupeneus barberinus* | 4 | Invertivore |
| *Plectroglyphidodon emeryi* | 1 | Corallivore, Herbivore |
| *Pomacanthus semicirculatus* | 1 | Herbivore, Invertivore |
| *Pomacentrus albicaudatus* | 51 | Invertivore |
| *Pomacentrus armillatus* | 1 | Herbivore |
| *Pomacentrus imitator* | 1 | Herbivore |
| *Pomacentrus leptus* | 2 | Herbivore, Invertivore |
| *Pomacentrus moluccensis* | 2 | Herbivore, Invertivore |
| *Pomacentrus nigromanus* | 1 | Herbivore, Invertivore |
| *Pomacentrus taeniometopon* | 2 | Herbivore, Invertivore |
| *Pomacentrus wardi* | 1 | Herbivore |
| *Scarus frenatus* | 28 | Herbivore |
| *Scarus ghobban* | 14 | Herbivore |
| *Scarus longipinnis* | 2 | Herbivore |
| *Scarus niger* | 28 | Herbivore |
| *Scarus quoyi* | 4 | Herbivore |
| *Scarus sordidus* | 1 | Herbivore |
| *Scolopsis affinis* | 5 | Invertivore, Piscivore |
| *Scolopsis bilineatus* | 8 | Invertivore, Piscivore |
| *Scolopsis margaritifera* | 5 | Invertivore, Piscivore |
| *Siganus javus* | 2 | Herbivore, Invertivore |
| *Stethojulis strigiventer* | 3 | Invertivore |
| *Stethojulis trilineata* | 4 | Invertivore |
| *Sufflamen chrysopterum* | 3 | Invertivore |
| *Thalassoma lunare* | 18 | Invertivore |
| *Variola louti* | 1 | Invertivore, Piscivore |
| *Zanclus cornutus* | 7 | Herbivore, Invertivore |
| *Zebrasoma desjardinii* | 1 | Herbivore |

## Appendix B: Summary of model for time budget

**Table 1:**
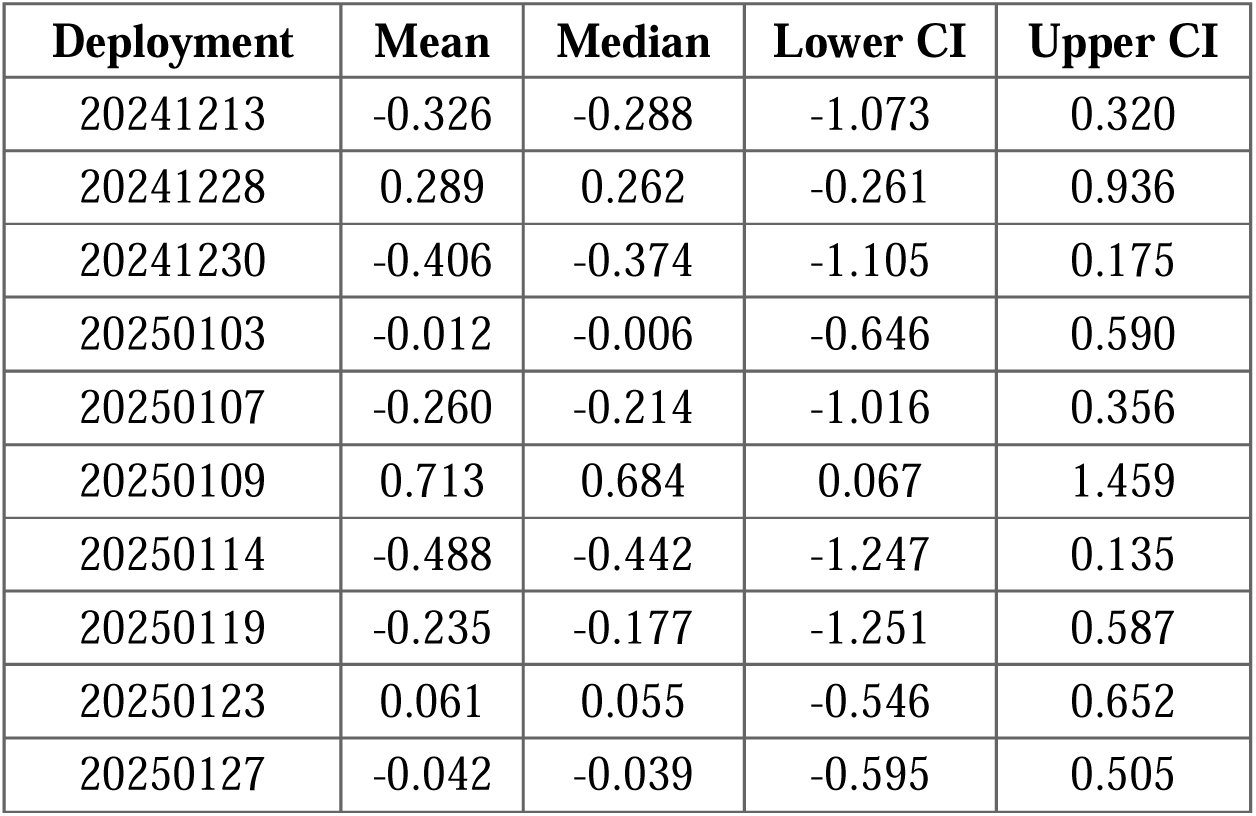
Summary of random intercepts for deployment for foraging time model.

**Table 2:** Summary of random intercepts of species for foraging time model.

| Family | Species | Mean | Median | Lower CI | Upper CI |
| --- | --- | --- | --- | --- | --- |
| Acanthuridae | *Acanthurus albipectoralis* | 1.196 | 1.165 | -0.168 | 2.671 |
|  | *Acanthurus blochii* | 0.327 | 0.331 | -0.538 | 1.189 |
|  | *Acanthurus leucosternon* | 0.703 | 0.635 | -0.782 | 2.430 |
|  | *Acanthurus lineatus* | -0.016 | -0.005 | -1.180 | 1.156 |
|  | *Acanthurus nigrofuscus* | 0.326 | 0.309 | -0.795 | 1.539 |
|  | *Ctenochaetus cyanocheilus* | -0.280 | -0.250 | -1.867 | 1.223 |
|  | *Ctenochaetus strigosus* | -0.473 | -0.429 | -1.970 | 0.871 |
|  | *Naso caesius* | -0.251 | -0.233 | -1.774 | 1.242 |
|  | *Naso elegans* | -0.543 | -0.487 | -2.049 | 0.772 |
|  | *Zebrasoma desjardini* | -0.245 | -0.213 | -1.828 | 1.197 |
| Balistidae | *Balistapus undulatus* | -0.339 | -0.294 | -1.867 | 1.050 |
|  | *Balistoides viridescens* | -0.257 | -0.241 | -1.841 | 1.233 |
|  | *Sufflamen chrysopterum* | 1.350 | 1.278 | -0.042 | 2.907 |
| Caesionidae | *Caesio caerulea* | -0.729 | -0.673 | -2.246 | 0.587 |
|  | *Caesio cuning* | -0.296 | -0.263 | -1.810 | 1.091 |
| Chaetodontidae | *Chaetodon andamanensis* | 0.433 | 0.408 | -0.374 | 1.313 |
|  | *Chaetodon auriga* | -0.108 | -0.089 | -1.701 | 1.434 |
|  | *Chaetodon baronessa* | 0.234 | 0.216 | -0.634 | 1.178 |
|  | *Chaetodon capistratus* | -0.252 | -0.208 | -1.781 | 1.196 |
|  | *Chaetodon decussatus* | -0.240 | -0.202 | -1.797 | 1.242 |
|  | *Chaetodon falcula* | 1.622 | 1.603 | 0.444 | 2.875 |
|  | *Chaetodon lunulatus* | 1.734 | 1.722 | 0.962 | 2.575 |
|  | *Chaetodon meyeri* | -0.438 | -0.383 | -2.089 | 0.953 |
|  | *Chaetodon triangulum* | -0.303 | -0.263 | -1.975 | 1.145 |
|  | *Chaetodon trifacialis* | 0.440 | 0.410 | -0.771 | 1.730 |
| Kyphosidae | *Kyphosus cinerascens* | -0.316 | -0.280 | -1.877 | 1.140 |
| Labridae | *Bodianus mesothorax* | -0.238 | -0.197 | -1.917 | 1.300 |
|  | *Cheilinus abudjubbe* | 0.563 | 0.532 | -0.357 | 1.605 |
|  | *Cheilinus chlorourus* | -0.349 | -0.307 | -1.971 | 1.176 |
|  | *Cheilinus fasciatus* | -0.152 | -0.120 | -1.694 | 1.326 |
|  | *Cheilinus indulatus* | -0.085 | -0.080 | -1.608 | 1.414 |
|  | *Cheilinus trilobatus* | -0.230 | -0.253 | -1.336 | 0.967 |
|  | *Epibulus insidiator* | -0.288 | -0.261 | -1.842 | 1.183 |
|  | *Gomphosus caeruleus* | -0.259 | -0.214 | -1.854 | 1.234 |
|  | *Halichoeres chierchiae* | 0.588 | 0.521 | -0.921 | 2.310 |
|  | *Halichoeres chloropterus* | -0.264 | -0.241 | -1.862 | 1.248 |
|  | *Halichoeres chrysotaenia* | 0.252 | 0.245 | -1.096 | 1.619 |
|  | *Halichoeres hortulanus* | 0.289 | 0.274 | -0.908 | 1.519 |
|  | *Halichoeres margaritaceus* | -0.217 | -0.177 | -1.905 | 1.285 |
|  | *Hemigymnus melapterus* | 0.660 | 0.641 | -0.439 | 1.823 |
|  | *Stethojulis strigiventer* | 0.710 | 0.623 | -0.764 | 2.412 |
|  | *Stethojulis trilineata* | -0.507 | -0.440 | -2.041 | 0.839 |
|  | *Thalassoma lunare* | -0.073 | -0.059 | -1.094 | 0.874 |
| Lethrinidae | *Lethrinus nebulosus* | -0.312 | -0.264 | -1.893 | 1.115 |
|  | *Lethrinus olivaceus* | -0.222 | -0.208 | -1.818 | 1.300 |
|  | *Monotaxis grandoculis* | -0.180 | -0.171 | -1.768 | 1.336 |
| Lutjanidae | *Lutjanus decussatus* | -1.024 | -0.962 | -2.515 | 0.221 |
|  | *Macolor niger* | -0.233 | -0.196 | -1.863 | 1.302 |
| Mullidae | *Parupeneus barberinus* | 0.227 | 0.228 | -1.140 | 1.580 |
| Nemipteridae | *Scolopsis affinis* | -0.739 | -0.668 | -2.179 | 0.518 |
|  | *Scolopsis bilineatus* | -0.628 | -0.553 | -2.093 | 0.618 |
|  | *Scolopsis margaritifera* | -0.734 | -0.669 | -2.305 | 0.545 |
| Pomacanthidae | *Centropyge eibli* | 1.117 | 1.093 | 0.151 | 2.157 |
|  | *Chaetodontoplus mesoleucus* | -0.085 | -0.069 | -1.384 | 1.217 |
|  | *Pomacanthus semicirculatus* | 0.696 | 0.625 | -0.796 | 2.454 |
| Pomacentridae | *Acanthochromis polyacanthus* | -0.295 | -0.267 | -1.842 | 1.172 |
|  | *Amblyglyphidodon aureus* | -0.342 | -0.297 | -1.911 | 1.058 |
|  | *Amblyglyphidodon batunaorum* | -0.239 | -0.231 | -1.850 | 1.253 |
|  | *Chromis jubauna* | 0.798 | 0.716 | -0.716 | 2.569 |
|  | *Chromis viridis* | -0.512 | -0.464 | -2.031 | 0.843 |
|  | *Chrysiptera talboti* | -0.487 | -0.441 | -2.055 | 0.918 |
|  | *Dascyllus reticulatus* | -0.641 | -0.592 | -2.207 | 0.744 |
|  | *Neoglyphidodon melas* | -0.407 | -0.364 | -2.018 | 0.977 |
|  | *Neopomacentrus nemurus* | -0.147 | -0.120 | -1.737 | 1.386 |
|  | *Plectroglyphidodon emeryi* | -0.134 | -0.124 | -1.781 | 1.465 |
|  | *Pomacentrus albicaudatus* | -0.078 | -0.083 | -0.711 | 0.574 |
|  | *Pomacentrus armillatus* | -0.273 | -0.227 | -1.941 | 1.199 |
|  | *Pomacentrus imitator* | -0.176 | -0.140 | -1.841 | 1.361 |
|  | *Pomacentrus leptus* | 0.399 | 0.383 | -0.689 | 1.540 |
|  | *Pomacentrus moluccensis* | -0.519 | -0.473 | -2.181 | 0.904 |
|  | *Pomacentrus nigromanus* | -0.027 | -0.041 | -1.148 | 1.148 |
|  | *Pomacentrus taeniometopon* | -0.351 | -0.315 | -1.892 | 1.054 |
| Scaridae | *Chlorurus sordidus* | -0.312 | -0.320 | -1.042 | 0.435 |
|  | *Scarus frenatus* | 0.574 | 0.562 | -0.536 | 1.731 |
|  | *Scarus ghobban* | -0.358 | -0.305 | -1.664 | 0.793 |
|  | *Scarus longipinnis* | -0.309 | -0.262 | -1.955 | 1.185 |
|  | *Scarus niger* | -0.268 | -0.269 | -1.152 | 0.618 |
|  | *Scarus quoyi* | 0.872 | 0.798 | -0.504 | 2.444 |
|  | *Scarus sordidus* | 0.546 | 0.502 | -0.901 | 2.165 |
| Serranidae | *Cephalopholis argus* | 0.025 | 0.036 | -1.252 | 1.296 |
|  | *Cephalopholis boenak* | -0.188 | -0.157 | -1.743 | 1.239 |
|  | *Cephalopholis formosa* | -0.232 | -0.210 | -1.757 | 1.267 |
|  | *Cephalopholis leopardus* | -0.109 | -0.087 | -1.679 | 1.424 |
|  | *Epinephelus fasciatus* | -0.258 | -0.220 | -1.813 | 1.201 |
|  | *Variola louti* | -0.166 | -0.128 | -1.736 | 1.327 |
| Siganidae | *Siganus javus* | -0.470 | -0.438 | -2.017 | 0.936 |
| Zanclidae | *Zanclus cornutus* | 0.601 | 0.582 | -0.373 | 1.599 |

**Table 3:** Summary of species-level random intercepts for precision for foraging time model.

| Family | Species | Mean | Median | Lower CI | Upper CI |
| --- | --- | --- | --- | --- | --- |
| Acanthuridae | *Acanthurus albipectoralis* | -1.458 | -1.377 | -3.685 | 0.258 |
|  | *Acanthurus blochii* | 0.533 | 0.352 | -1.273 | 2.908 |
|  | *Acanthurus leucosternon* | 0.031 | -0.003 | -2.375 | 2.648 |
|  | *Acanthurus lineatus* | 0.031 | 0.000 | -2.371 | 2.398 |
|  | *Acanthurus nigrofuscus* | 0.019 | -0.007 | -2.438 | 2.574 |
|  | *Ctenochaetus cyanocheilus* | -0.032 | 0.012 | -2.572 | 2.401 |
|  | *Ctenochaetus strigosus* | 0.051 | 0.015 | -2.445 | 2.570 |
|  | *Naso caesius* | 0.013 | -0.001 | -2.376 | 2.460 |
|  | *Naso elegans* | 0.009 | 0.004 | -2.428 | 2.341 |
|  | *Zebbrasoma desjardinii* | 0.017 | 0.001 | -2.307 | 2.449 |
| Balistidae | *Balistapus undulatus* | 0.022 | -0.003 | -2.442 | 2.466 |
|  | *Balistoides viridescens* | -0.001 | 0.013 | -2.536 | 2.369 |
|  | *Sufflamen chrysopterum* | 0.068 | 0.027 | -2.345 | 2.638 |
| Caesionidae | *Caesio caeruleaurea* | -0.019 | -0.024 | -2.386 | 2.408 |
|  | *Caesio cuning* | 0.044 | 0.036 | -2.475 | 2.451 |
| Chaetodontidae | *Chaetodon andamanensis* | -0.087 | -0.085 | -1.935 | 1.870 |
|  | *Chaetodon auriga* | 0.012 | -0.013 | -2.516 | 2.528 |
|  | *Chaetodon baronessa* | 0.361 | 0.201 | -1.618 | 2.779 |
|  | *Chaetodon capistratus* | -0.015 | 0.004 | -2.482 | 2.542 |
|  | *Chaetodon decussatus* | -0.015 | -0.015 | -2.505 | 2.458 |
|  | *Chaetodon falcula* | 0.311 | 0.152 | -1.617 | 2.722 |
|  | *Chaetodon lunulatus* | 1.368 | 1.286 | -0.273 | 3.405 |
|  | *Chaetodon meyeri* | -0.001 | 0.000 | -2.494 | 2.419 |
|  | *Chaetodon triangulum* | 0.034 | 0.006 | -2.432 | 2.580 |
|  | *Chaetodon trifacialis* | -0.048 | -0.020 | -2.566 | 2.384 |
| Kyphosidae | *Kyphosus cinerascens* | -0.024 | -0.013 | -2.440 | 2.447 |
| Labridae | *Bodianus mesothorax* | 0.041 | 0.029 | -2.351 | 2.423 |
|  | *Cheilinus abudjubbe* | 0.324 | 0.200 | -1.730 | 2.634 |
|  | *Cheilinus chlorourus* | 0.008 | -0.001 | -2.398 | 2.495 |
|  | *Cheilinus fasciatus* | 0.021 | 0.002 | -2.363 | 2.636 |
|  | *Cheilinus indulatus* | -0.005 | 0.008 | -2.400 | 2.413 |
|  | *Cheilinus trilobatus* | -0.286 | -0.261 | -2.497 | 1.982 |
|  | *Epibulus insidiator* | -0.014 | -0.011 | -2.465 | 2.503 |
|  | *Gomphosus caeruleus* | 0.028 | 0.008 | -2.423 | 2.451 |
|  | *Halichoeres chierchiae* | -0.039 | 0.003 | -2.588 | 2.345 |
|  | *Halichoeres chloropterus* | 0.002 | 0.009 | -2.398 | 2.445 |
|  | *Halichoeres chrysotaenia* | 0.004 | -0.004 | -2.474 | 2.479 |
|  | *Halichoeres hortulanus* | -0.014 | -0.002 | -2.412 | 2.411 |
|  | *Halichoeres margaritaceus* | 0.012 | 0.003 | -2.536 | 2.455 |
|  | *Hemigymnus melapterus* | 0.378 | 0.237 | -1.539 | 2.677 |
|  | *Stethojulis strigiventer* | 0.016 | -0.007 | -2.483 | 2.676 |
|  | *Stethojulis trilineata* | -0.010 | 0.008 | -2.533 | 2.390 |
|  | *Thalassoma lunare* | 0.342 | 0.218 | -1.705 | 2.651 |
| Lethrinidae | *Lethrinus nebulosus* | 0.003 | 0.001 | -2.606 | 2.540 |
|  | *Lethrinus olivaceus* | 0.033 | -0.001 | -2.316 | 2.388 |
|  | *Monotaxis grandoculis* | 0.033 | 0.019 | -2.233 | 2.486 |
| Lutjanidae | *Lutjanus decussatus* | 0.039 | 0.008 | -2.467 | 2.584 |
|  | *Macolor niger* | -0.015 | -0.022 | -2.508 | 2.462 |
| Mullidae | *Parupeneus barberinus* | 0.037 | 0.020 | -2.455 | 2.592 |
| Nemipteridae | *Scolopsis affinis* | 0.004 | 0.021 | -2.344 | 2.387 |
|  | *Scolopsis bilineatus* | -0.021 | -0.007 | -2.666 | 2.615 |
|  | *Scolopsis margaritifera* | 0.030 | 0.010 | -2.310 | 2.524 |
| Pomacanthidae | *Centropyge eibli* | -1.231 | -1.132 | -3.237 | 0.301 |
|  | *Chaetodontoplus mesoleucus* | 0.015 | 0.009 | -2.441 | 2.473 |
|  | *Pomacanthus semicirculatus* | 0.008 | -0.007 | -2.390 | 2.562 |
| Pomacentridae | *Acanthochromis polyacanthus* | 0.007 | -0.001 | -2.410 | 2.478 |
|  | *Amblyglyphidodon aureus* | -0.015 | 0.000 | -2.642 | 2.557 |
|  | *Amblyglyphidodon batunaorum* | 0.015 | -0.008 | -2.427 | 2.543 |
|  | *Chromis jubauna* | -0.002 | -0.010 | -2.386 | 2.445 |
|  | *Chromis viridis* | -0.007 | -0.011 | -2.455 | 2.630 |
|  | *Chrysiptera talboti* | -0.062 | -0.007 | -2.676 | 2.398 |
|  | *Dascyllus reticulatus* | 0.040 | 0.005 | -2.337 | 2.471 |
|  | *Neoglyphidodon melas* | -0.049 | -0.032 | -2.399 | 2.275 |
|  | *Neopomacentrus nemurus* | -0.012 | -0.010 | -2.377 | 2.420 |
|  | *Plectroglyphidodon emeryi* | 0.044 | 0.007 | -2.377 | 2.601 |
|  | *Pomacentrus albicaudatus* | -0.883 | -0.816 | -2.258 | 0.236 |
|  | *Pomacentrus armillatus* | -0.047 | -0.005 | -2.461 | 2.283 |
|  | *Pomacentrus imitator* | -0.016 | 0.000 | -2.623 | 2.484 |
|  | *Pomacentrus leptus* | 0.369 | 0.208 | -1.611 | 2.698 |
|  | *Pomacentrus moluccensis* | -0.011 | -0.026 | -2.455 | 2.523 |
|  | *Pomacentrus nigromanus* | 0.180 | 0.078 | -2.008 | 2.746 |
|  | *Pomacentrus taeniometopon* | 0.009 | -0.006 | -2.368 | 2.446 |
| Scaridae | *Chlorurus sordidus* | 0.393 | 0.271 | -1.148 | 2.319 |
|  | *Scarus frenatus* | 0.041 | 0.004 | -2.515 | 2.649 |
|  | *Scarus ghobban* | 0.011 | 0.006 | -2.420 | 2.514 |
|  | *Scarus longipinnis* | 0.023 | 0.005 | -2.260 | 2.310 |
|  | *Scarus niger* | 0.262 | 0.128 | -1.782 | 2.670 |
|  | *Scarus quoyi* | -0.039 | 0.000 | -2.560 | 2.319 |
|  | *Scarus sordidus* | -0.012 | 0.002 | -2.498 | 2.382 |
| Serranidae | *Cephalopholis argus* | -0.005 | 0.006 | -2.548 | 2.437 |
|  | *Cephalopholis boenak* | -0.017 | 0.004 | -2.631 | 2.433 |
|  | *Cephalopholis formosa* | 0.033 | -0.002 | -2.202 | 2.420 |
|  | *Cephalopholis leopardus* | -0.050 | -0.012 | -2.725 | 2.436 |
|  | *Epinephelus fasciatus* | -0.015 | -0.010 | -2.433 | 2.272 |
|  | *Variola louti* | 0.001 | -0.010 | -2.413 | 2.392 |
| Siganidae | *Siganus javus* | 0.014 | 0.009 | -2.295 | 2.338 |
| Zanclidae | *Zanclus cornutus* | 0.116 | 0.118 | -1.575 | 1.672 |

**Table 4:** Summary of hyperparameters for foraging time mode.

| Variable | Mean | Median | Lower CI | Upper CI | P(>0) |
| --- | --- | --- | --- | --- | --- |
| Cutone | -0.345 | -0.341 | -0.628 | -0.076 | 0.016 |
| Cutzero | 0.510 | 0.508 | 0.162 | 0.865 | 0.993 |

**Table 5:** Summary of fixed effects for foraging time model.

| **Variable** | **Mean** | **Median** | **Lower CI** | **Upper CI** | **P(>0)** |
| --- | --- | --- | --- | --- | --- |
| Intercept | -0.458 | -0.469 | -1.315 | 0.443 | 0.196 |
| Protection | 0.137 | 0.146 | -0.519 | 0.780 | 0.640 |
| Treatment (Grouper) | -0.282 | -0.279 | -0.919 | 0.325 | 0.226 |
| Treatment (Negative Control) | 0.145 | 0.145 | -0.456 | 0.750 | 0.649 |
| Treatment (Barracuda) | -0.062 | -0.060 | -0.686 | 0.548 | 0.435 |
| Guild (Invertivore) | -0.053 | -0.049 | -0.723 | 0.612 | 0.447 |
| Rugosity | -0.328 | -0.330 | -0.589 | -0.052 | 0.025 |
| Biomass | -0.068 | -0.065 | -0.314 | 0.172 | 0.325 |
| Grouping | -0.067 | -0.090 | -0.388 | 0.331 | 0.343 |
| Size class: 10-20 | 0.285 | 0.282 | -0.171 | 0.741 | 0.850 |
| Size class: 20-30 | -0.042 | -0.047 | -0.562 | 0.493 | 0.438 |
| Size class: 30-40 | 0.017 | 0.013 | -0.731 | 0.799 | 0.512 |
| Size class: 40-50 | 0.301 | 0.298 | -0.448 | 1.049 | 0.746 |
| Protection:Treatment (Grouper) | 0.150 | 0.148 | -0.499 | 0.808 | 0.649 |
| Protection:Treatment (Negative Control) | -0.300 | -0.294 | -0.971 | 0.366 | 0.242 |
| Protection:Treatment (Barracuda) | -0.191 | -0.186 | -0.848 | 0.469 | 0.316 |
| Treatment (Grouper):Guild (Invertivore) | -0.122 | -0.113 | -0.766 | 0.505 | 0.372 |
| Treatment (Negative Control):Guild (Invertivore) | 0.089 | 0.089 | -0.564 | 0.748 | 0.583 |
| Treatment (Barracuda):Guild (Invertivore) | 0.018 | 0.021 | -0.613 | 0.646 | 0.523 |
| Protection:Treatment (Negative Control):Guild (Invertivore) | -0.149 | -0.149 | -0.857 | 0.566 | 0.368 |
| Protection:Treatment (Grouper):Guild (Invertivore) | 0.284 | 0.292 | -0.389 | 0.940 | 0.761 |
| Protection:Treatment (Barracuda):Guild (Invertivore) | -0.346 | -0.345 | -1.044 | 0.322 | 0.206 |

**Table 6:**
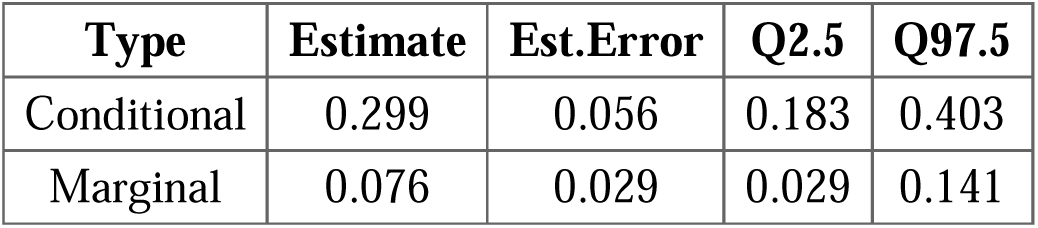
Summary of model fit for foraging time model.

| **Type** | **Estimate** | **Est.Error** | **Q2.5** | **Q97.5** |
| --- | --- | --- | --- | --- |
| Conditional | 0.299 | 0.056 | 0.183 | 0.403 |
| Marginal | 0.076 | 0.029 | 0.029 | 0.141 |

**Table 7:** Summary of random intercepts for deployment for vigilance time model.

| **Deployment** | **Mean** | **Median** | **Lower CI** | **Upper CI** |
| --- | --- | --- | --- | --- |
| 20241213 | -0.327 | -0.237 | -1.053 | 0.092 |
| 20241228 | -0.124 | -0.057 | -0.705 | 0.294 |
| 20241230 | 0.169 | 0.109 | -0.253 | 0.731 |
| 20250103 | 0.142 | 0.105 | -0.271 | 0.668 |
| 20250107 | 0.070 | 0.036 | -0.414 | 0.620 |
| 20250109 | -0.069 | -0.023 | -0.616 | 0.385 |
| 20250114 | 0.024 | 0.013 | -0.457 | 0.518 |
| 20250119 | -0.300 | -0.144 | -1.330 | 0.256 |
| 20250123 | -0.061 | -0.020 | -0.611 | 0.383 |
| 20250127 | -0.294 | -0.211 | -1.007 | 0.131 |

**Table 8:**
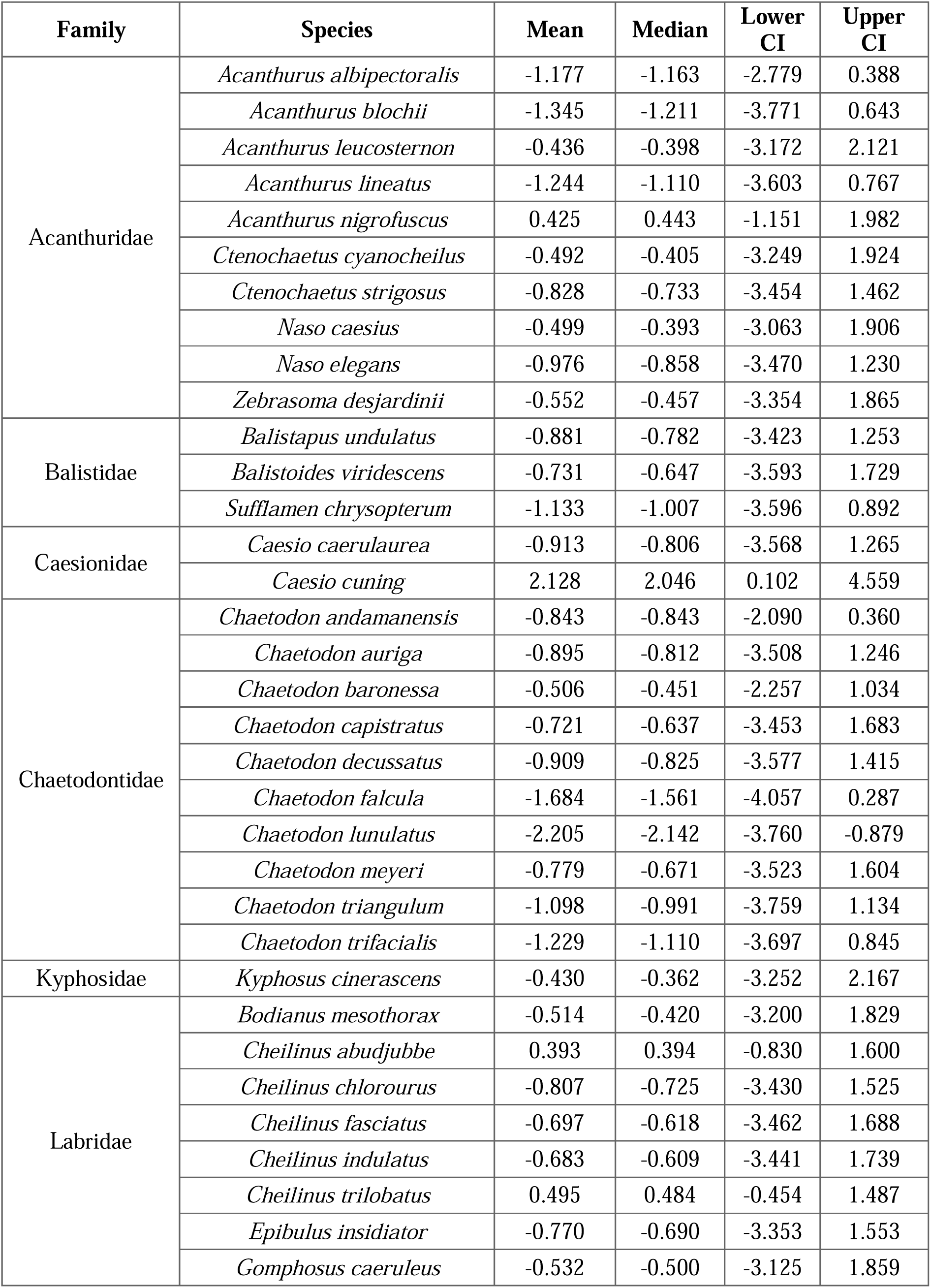
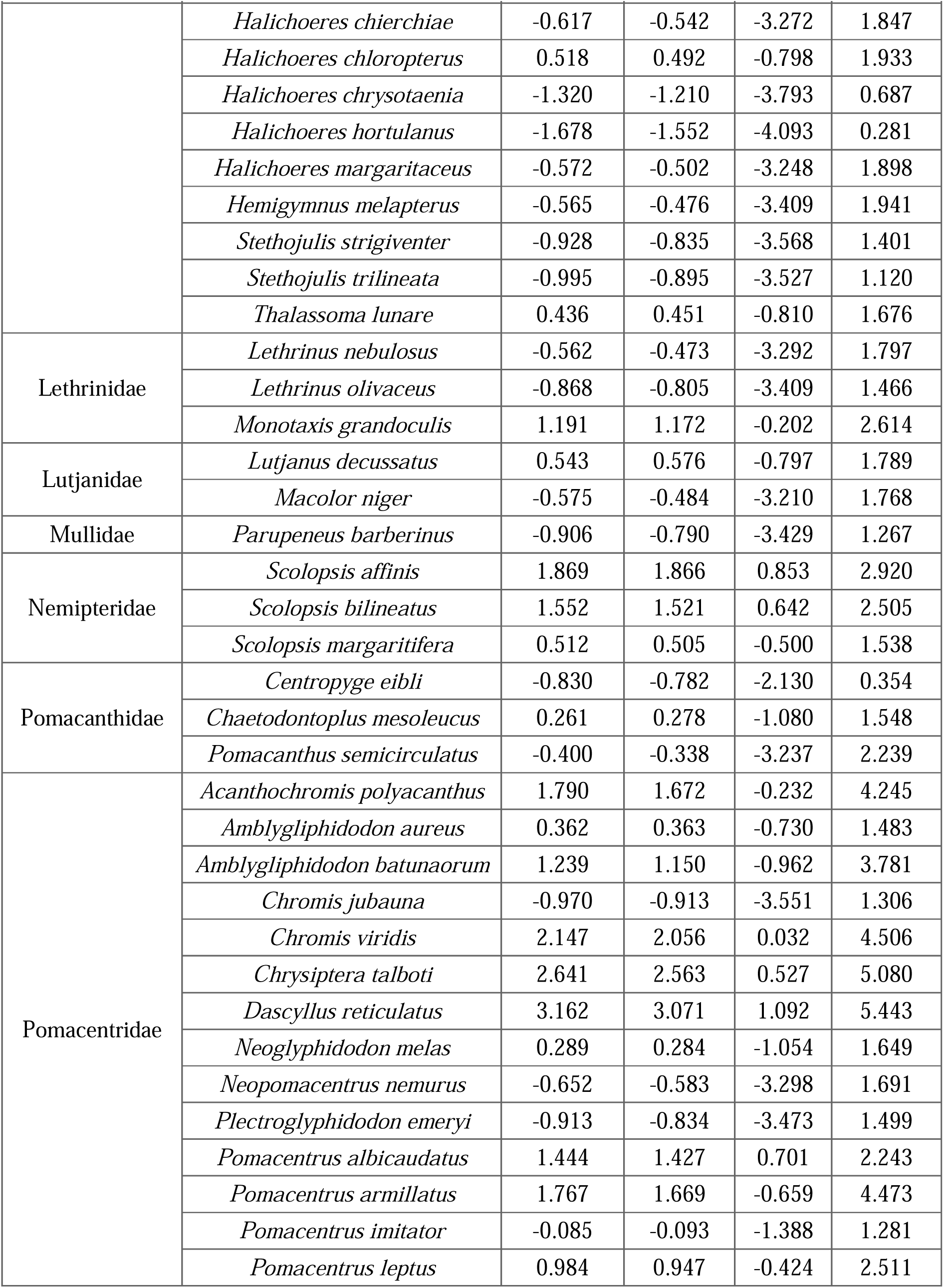
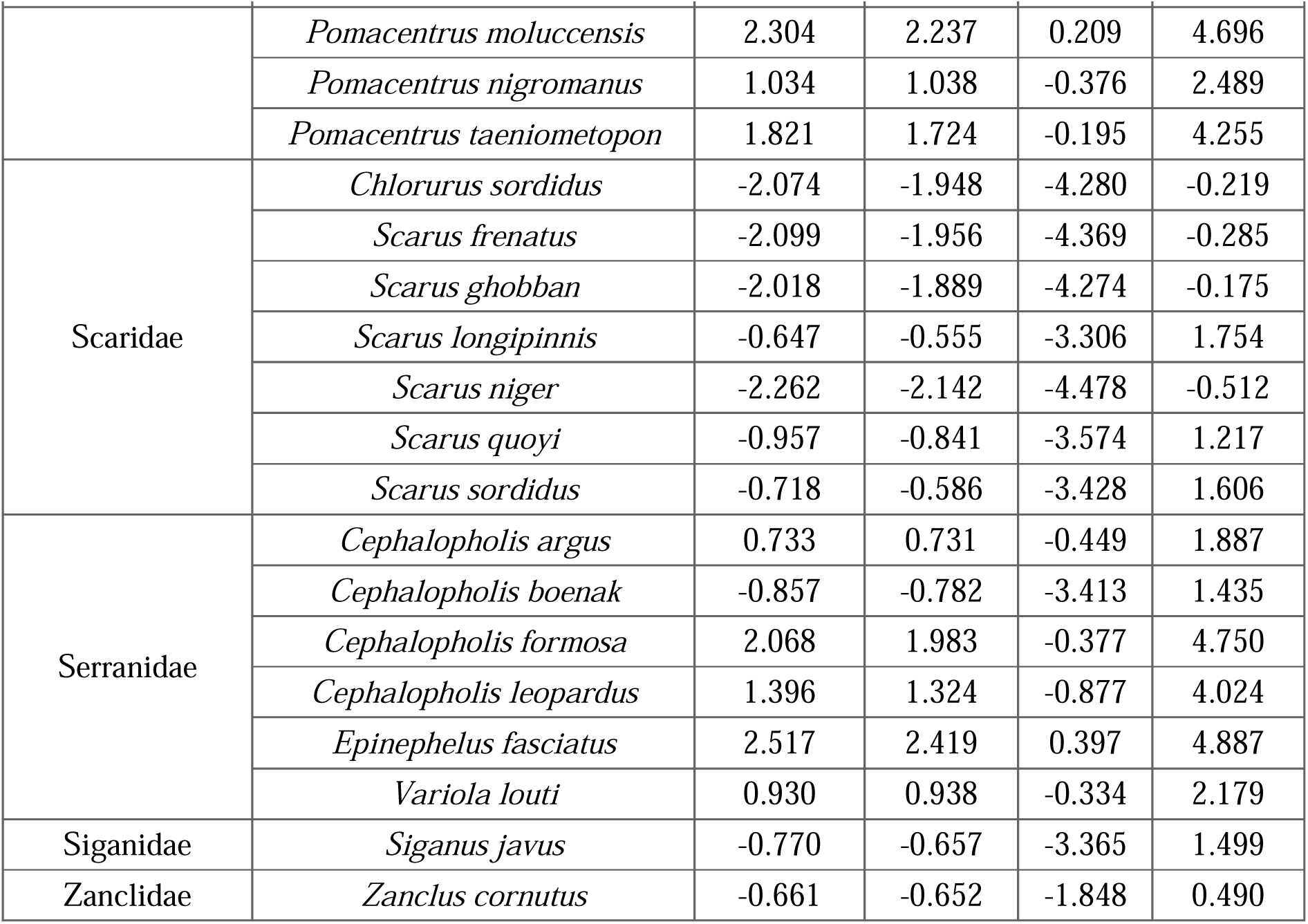
Summary of random intercepts of species for vigilance time model.

**Table 9:**
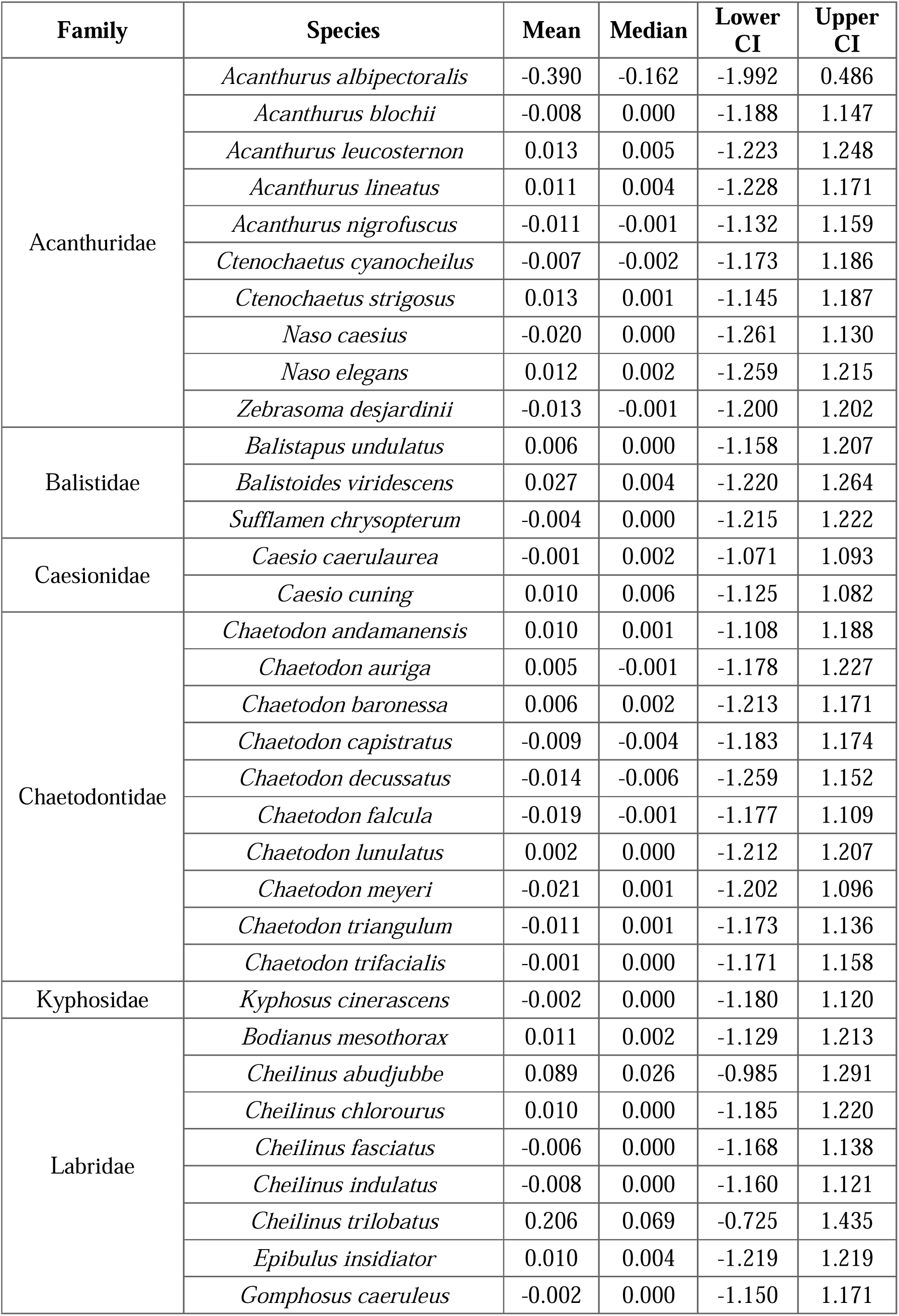
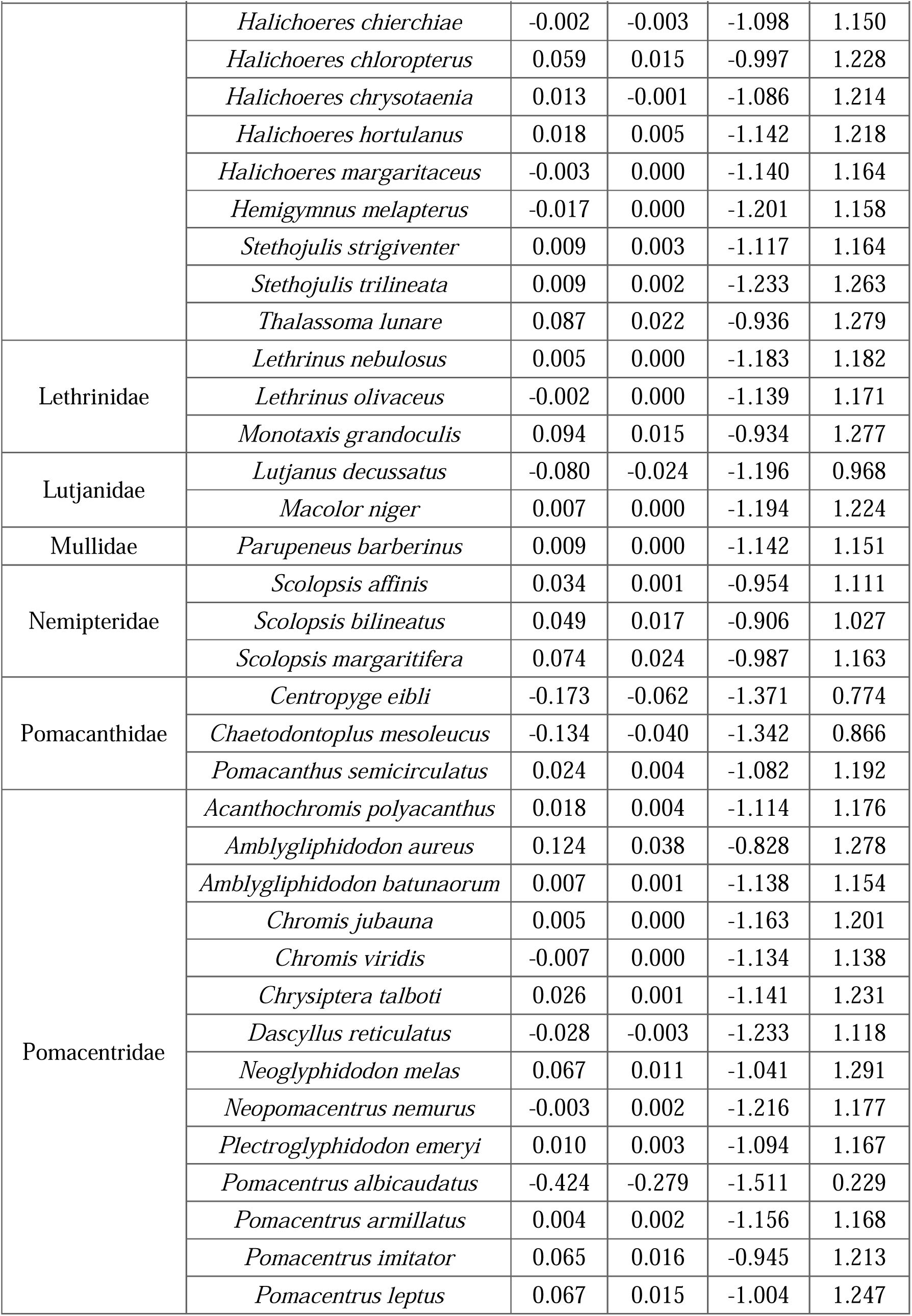
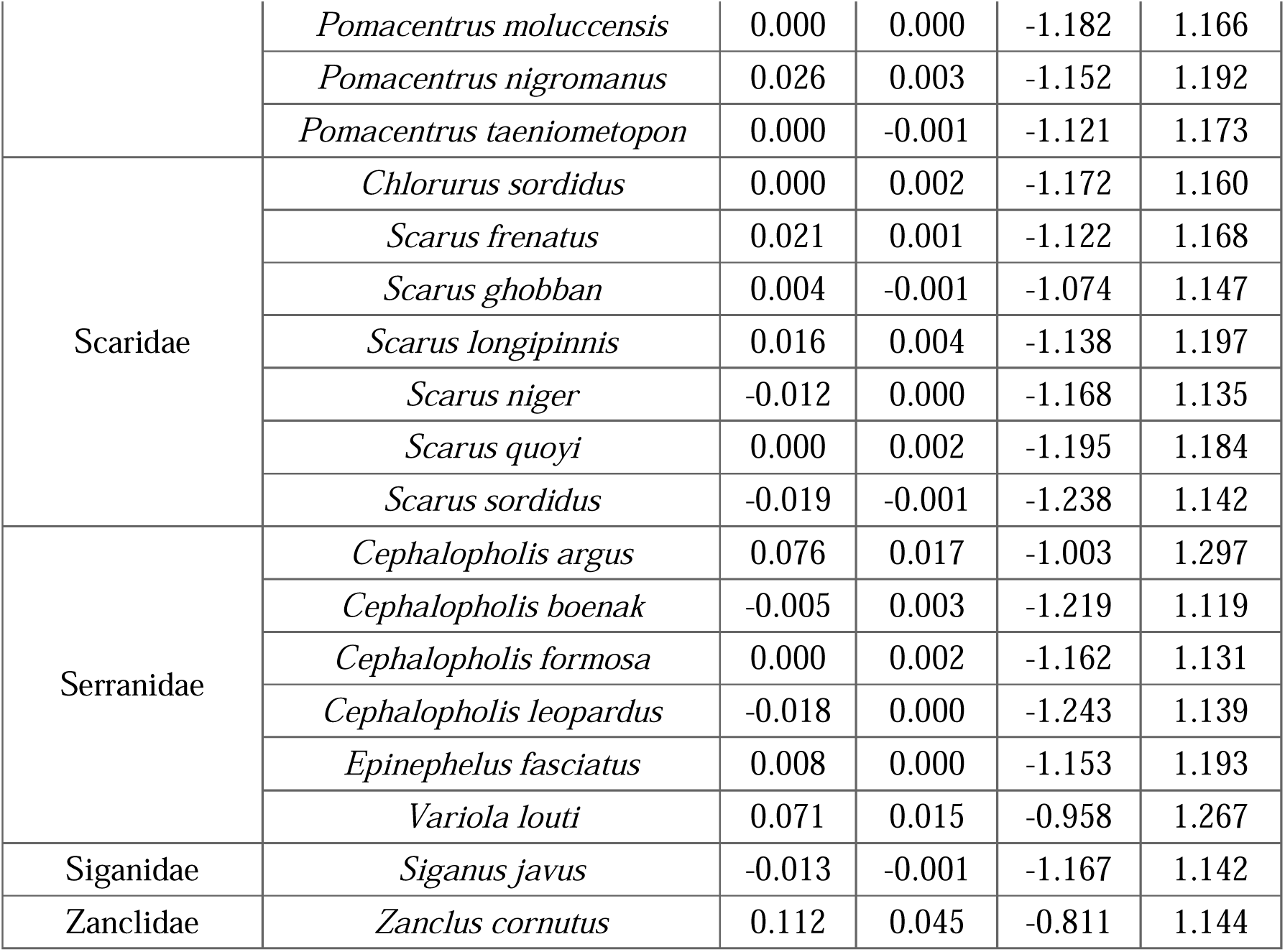
Summary of species-level random intercepts for precision for vigilance time model.

| Family | Species | Mean | Median | Lower CI | Upper CI |
| --- | --- | --- | --- | --- | --- |
| Acanthuridae | *Acanthurus albipectoralis* | -0.390 | -0.162 | -1.992 | 0.486 |
|  | *Acanthurus blochii* | -0.008 | 0.000 | -1.188 | 1.147 |
|  | *Acanthurus leucosternon* | 0.013 | 0.005 | -1.223 | 1.248 |
|  | *Acanthurus lineatus* | 0.011 | 0.004 | -1.228 | 1.171 |
|  | *Acanthurus nigrofuscus* | -0.011 | -0.001 | -1.132 | 1.159 |
|  | *Ctenochaetus cyanocheilus* | -0.007 | -0.002 | -1.173 | 1.186 |
|  | *Ctenochaetus strigosus* | 0.013 | 0.001 | -1.145 | 1.187 |
|  | *Naso caesius* | -0.020 | 0.000 | -1.261 | 1.130 |
|  | *Naso elegans* | 0.012 | 0.002 | -1.259 | 1.215 |
|  | *Zebrasoma desjardini* | -0.013 | -0.001 | -1.200 | 1.202 |
| Balistidae | *Balistapus undulatus* | 0.006 | 0.000 | -1.158 | 1.207 |
|  | *Balistoides viridescens* | 0.027 | 0.004 | -1.220 | 1.264 |
|  | *Sufflamen chrysopterum* | -0.004 | 0.000 | -1.215 | 1.222 |
| Caesionidae | *Caesio caerulaurea* | -0.001 | 0.002 | -1.071 | 1.093 |
|  | *Caesio cuning* | 0.010 | 0.006 | -1.125 | 1.082 |
| Chaetodontidae | *Chaetodon andamanensis* | 0.010 | 0.001 | -1.108 | 1.188 |
|  | *Chaetodon auriga* | 0.005 | -0.001 | -1.178 | 1.227 |
|  | *Chaetodon baronessa* | 0.006 | 0.002 | -1.213 | 1.171 |
|  | *Chaetodon capistratus* | -0.009 | -0.004 | -1.183 | 1.174 |
|  | *Chaetodon decussatus* | -0.014 | -0.006 | -1.259 | 1.152 |
|  | *Chaetodon falcula* | -0.019 | -0.001 | -1.177 | 1.109 |
|  | *Chaetodon lunulatus* | 0.002 | 0.000 | -1.212 | 1.207 |
|  | *Chaetodon meyeri* | -0.021 | 0.001 | -1.202 | 1.096 |
|  | *Chaetodon triangulum* | -0.011 | 0.001 | -1.173 | 1.136 |
|  | *Chaetodon trifacialis* | -0.001 | 0.000 | -1.171 | 1.158 |
| Kyphosidae | *Kyphosus cinerascens* | -0.002 | 0.000 | -1.180 | 1.120 |
| Labridae | *Bodianus mesothorax* | 0.011 | 0.002 | -1.129 | 1.213 |
|  | *Cheilinus abudjubbe* | 0.089 | 0.026 | -0.985 | 1.291 |
|  | *Cheilinus chlorourus* | 0.010 | 0.000 | -1.185 | 1.220 |
|  | *Cheilinus fasciatus* | -0.006 | 0.000 | -1.168 | 1.138 |
|  | *Cheilinus indulatus* | -0.008 | 0.000 | -1.160 | 1.121 |
|  | *Cheilinus trilobatus* | 0.206 | 0.069 | -0.725 | 1.435 |
|  | *Epibulus insidiator* | 0.010 | 0.004 | -1.219 | 1.219 |
|  | *Gomphosus caeruleus* | -0.002 | 0.000 | -1.150 | 1.171 |
|  | *Halichoeres chierchiae* | -0.002 | -0.003 | -1.098 | 1.150 |
|  | *Halichoeres chloropterus* | 0.059 | 0.015 | -0.997 | 1.228 |
|  | *Halichoeres chrysotaenia* | 0.013 | -0.001 | -1.086 | 1.214 |
|  | *Halichoeres hortulanus* | 0.018 | 0.005 | -1.142 | 1.218 |
|  | *Halichoeres margaritaceus* | -0.003 | 0.000 | -1.140 | 1.164 |
|  | *Hemigymnus melapterus* | -0.017 | 0.000 | -1.201 | 1.158 |
|  | *Stethojulis strigiventer* | 0.009 | 0.003 | -1.117 | 1.164 |
|  | *Stethojulis trilineata* | 0.009 | 0.002 | -1.233 | 1.263 |
|  | *Thalassoma lunare* | 0.087 | 0.022 | -0.936 | 1.279 |
| Lethrinidae | *Lethrinus nebulosus* | 0.005 | 0.000 | -1.183 | 1.182 |
|  | *Lethrinus olivaceus* | -0.002 | 0.000 | -1.139 | 1.171 |
|  | *Monotaxis grandoculis* | 0.094 | 0.015 | -0.934 | 1.277 |
| Lutjanidae | *Lutjanus decussatus* | -0.080 | -0.024 | -1.196 | 0.968 |
|  | *Macolor niger* | 0.007 | 0.000 | -1.194 | 1.224 |
| Mullidae | *Parupeneus barberinus* | 0.009 | 0.000 | -1.142 | 1.151 |
| Nemipteridae | *Scolopsis affinis* | 0.034 | 0.001 | -0.954 | 1.111 |
|  | *Scolopsis bilineatus* | 0.049 | 0.017 | -0.906 | 1.027 |
|  | *Scolopsis margaritifera* | 0.074 | 0.024 | -0.987 | 1.163 |
| Pomacanthidae | *Centropyge eibli* | -0.173 | -0.062 | -1.371 | 0.774 |
|  | *Chaetodontoplus mesoleucus* | -0.134 | -0.040 | -1.342 | 0.866 |
|  | *Pomacanthus semicirculatus* | 0.024 | 0.004 | -1.082 | 1.192 |
| Pomacentridae | *Acanthochromis polyacanthus* | 0.018 | 0.004 | -1.114 | 1.176 |
|  | *Amblyglyphidodon aureus* | 0.124 | 0.038 | -0.828 | 1.278 |
|  | *Amblyglyphidodon batunaorum* | 0.007 | 0.001 | -1.138 | 1.154 |
|  | *Chromis jubauna* | 0.005 | 0.000 | -1.163 | 1.201 |
|  | *Chromis viridis* | -0.007 | 0.000 | -1.134 | 1.138 |
|  | *Chrysiptera talboti* | 0.026 | 0.001 | -1.141 | 1.231 |
|  | *Dascyllus reticulatus* | -0.028 | -0.003 | -1.233 | 1.118 |
|  | *Neoglyphidodon melas* | 0.067 | 0.011 | -1.041 | 1.291 |
|  | *Neopomacentrus nemurus* | -0.003 | 0.002 | -1.216 | 1.177 |
|  | *Plectroglyphidodon emeryi* | 0.010 | 0.003 | -1.094 | 1.167 |
|  | *Pomacentrus albicaudatus* | -0.424 | -0.279 | -1.511 | 0.229 |
|  | *Pomacentrus armillatus* | 0.004 | 0.002 | -1.156 | 1.168 |
|  | *Pomacentrus imitator* | 0.065 | 0.016 | -0.945 | 1.213 |
|  | *Pomacentrus leptus* | 0.067 | 0.015 | -1.004 | 1.247 |
|  | *Pomacentrus moluccensis* | 0.000 | 0.000 | -1.182 | 1.166 |
|  | *Pomacentrus nigromanus* | 0.026 | 0.003 | -1.152 | 1.192 |
|  | *Pomacentrus taeniometopon* | 0.000 | -0.001 | -1.121 | 1.173 |
| Scaridae | *Chlorurus sordidus* | 0.000 | 0.002 | -1.172 | 1.160 |
|  | *Scarus frenatus* | 0.021 | 0.001 | -1.122 | 1.168 |
|  | *Scarus ghobban* | 0.004 | -0.001 | -1.074 | 1.147 |
|  | *Scarus longipinnis* | 0.016 | 0.004 | -1.138 | 1.197 |
|  | *Scarus niger* | -0.012 | 0.000 | -1.168 | 1.135 |
|  | *Scarus quoyi* | 0.000 | 0.002 | -1.195 | 1.184 |
|  | *Scarus sordidus* | -0.019 | -0.001 | -1.238 | 1.142 |
| Serranidae | *Cephalopholis argus* | 0.076 | 0.017 | -1.003 | 1.297 |
|  | *Cephalopholis boenak* | -0.005 | 0.003 | -1.219 | 1.119 |
|  | *Cephalopholis formosa* | 0.000 | 0.002 | -1.162 | 1.131 |
|  | *Cephalopholis leopardus* | -0.018 | 0.000 | -1.243 | 1.139 |
|  | *Epinephelus fasciatus* | 0.008 | 0.000 | -1.153 | 1.193 |
|  | *Variola louti* | 0.071 | 0.015 | -0.958 | 1.267 |
| Siganidae | *Siganus javus* | -0.013 | -0.001 | -1.167 | 1.142 |
| Zanclidae | *Zanclus cornutus* | 0.112 | 0.045 | -0.811 | 1.144 |

**Table 10:** Summary of hyperparameters for vigilance time mode.

| Variable | Mean | Median | Lower CI | Upper CI | P(>0) |
| --- | --- | --- | --- | --- | --- |
| Cutone | 0.137 | 0.138 | -0.113 | 0.383 | 0.827 |
| Cutzero | -0.061 | -0.061 | -0.436 | 0.324 | 0.397 |

**Table 11:** Summary of fixed effects for vigilance time model.

| **Variable** | **Mean** | **Median** | **Lower CI** | **Upper CI** | **P(>0)** |
| --- | --- | --- | --- | --- | --- |
| Intercept | -1.377 | -1.376 | -2.386 | -0.401 | 0.013 |
| Protection | 0.052 | 0.047 | -0.583 | 0.668 | 0.550 |
| Treatment (Grouper) | 0.147 | 0.140 | -0.449 | 0.757 | 0.647 |
| Treatment (Negative Control) | 0.300 | 0.303 | -0.333 | 0.926 | 0.790 |
| Treatment (Barracuda) | 0.172 | 0.170 | -0.471 | 0.808 | 0.671 |
| Guild (Invertivore) | -0.009 | -0.009 | -0.714 | 0.718 | 0.490 |
| Rugosity | 0.080 | 0.080 | -0.148 | 0.315 | 0.720 |
| Biomass | -0.005 | -0.005 | -0.230 | 0.220 | 0.485 |
| Group | -0.205 | -0.206 | -0.811 | 0.392 | 0.286 |
| Size class: 10-20 | -0.051 | -0.055 | -0.598 | 0.515 | 0.435 |
| Size class: 20-30 | -1.004 | -1.008 | -1.645 | -0.356 | 0.005 |
| Size class: 30-40 | -0.105 | -0.115 | -0.897 | 0.712 | 0.414 |
| Size class: 40-50 | -0.013 | -0.016 | -0.824 | 0.809 | 0.484 |
| Protection:Treatment (Grouper) | 0.438 | 0.445 | -0.208 | 1.071 | 0.875 |
| Protection:Treatment (Negative Control) | 0.115 | 0.110 | -0.569 | 0.799 | 0.606 |
| Protection:Treatment (Barracuda) | 0.510 | 0.506 | -0.141 | 1.177 | 0.903 |
| Treatment (Grouper):Guild (Invertivore) | 0.233 | 0.236 | -0.365 | 0.844 | 0.731 |
| Treatment (Negative Control):Guild (Invertivore) | 0.280 | 0.283 | -0.363 | 0.900 | 0.772 |
| Treatment (Barracuda):Guild (Invertivore) | 0.064 | 0.067 | -0.586 | 0.701 | 0.571 |
| Protection:Treatment (Negative Control):Guild (Invertivore) | 0.092 | 0.099 | -0.639 | 0.795 | 0.588 |
| Protection:Treatment (Grouper):Guild (Invertivore) | 0.123 | 0.121 | -0.549 | 0.789 | 0.617 |
| Protection:Treatment (Barracuda):Guild (Invertivore) | 0.339 | 0.343 | -0.360 | 1.023 | 0.795 |

**Table 12:** Summary of model fit for vigilance time model.

| **Type** | **Estimate** | **Est.Error** | **Q2.5** | **Q97.5** |
| --- | --- | --- | --- | --- |
| Conditional | 0.484 | 0.056 | 0.370 | 0.587 |
| Marginal | 0.115 | 0.042 | 0.042 | 0.203 |

**Table 13:** Summary of random intercepts for deployment for movement time model.

| **Deployment** | **Mean** | **Median** | **Lower CI** | **Upper CI** |
| --- | --- | --- | --- | --- |
| 20241213 | -0.368 | -0.297 | -1.249 | 0.276 |
| 20241228 | -0.294 | -0.258 | -0.942 | 0.238 |
| 20241230 | 0.122 | 0.085 | -0.417 | 0.794 |
| 20250103 | 0.078 | 0.059 | -0.481 | 0.667 |
| 20250107 | 0.108 | 0.084 | -0.429 | 0.695 |
| 20250109 | -0.580 | -0.539 | -1.289 | -0.014 |
| 20250114 | 0.373 | 0.322 | -0.156 | 1.035 |
| 20250119 | -0.135 | -0.081 | -0.969 | 0.601 |
| 20250123 | 0.132 | 0.110 | -0.482 | 0.828 |
| 20250127 | 0.360 | 0.340 | -0.131 | 0.960 |

**Table 14:**
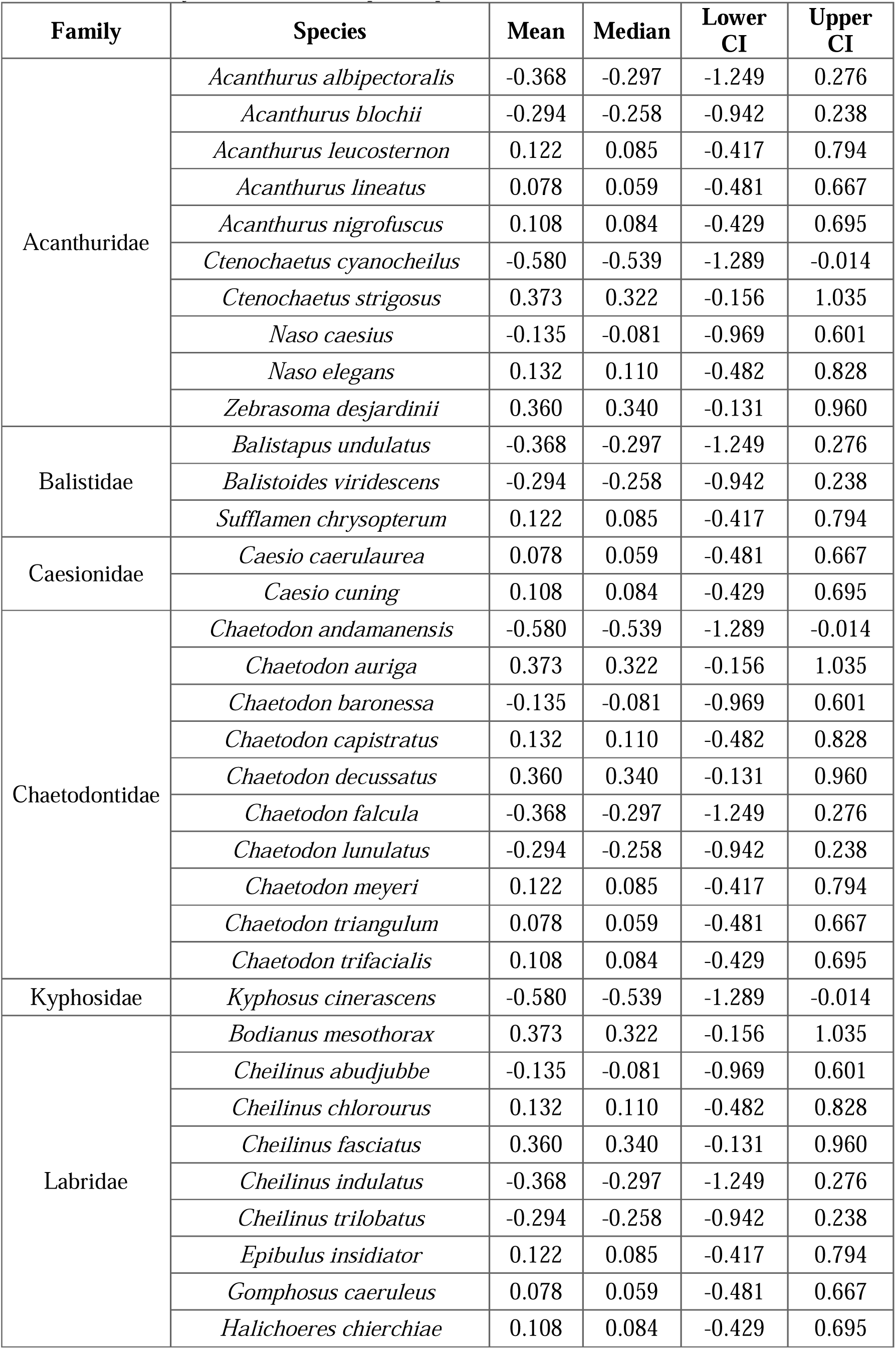
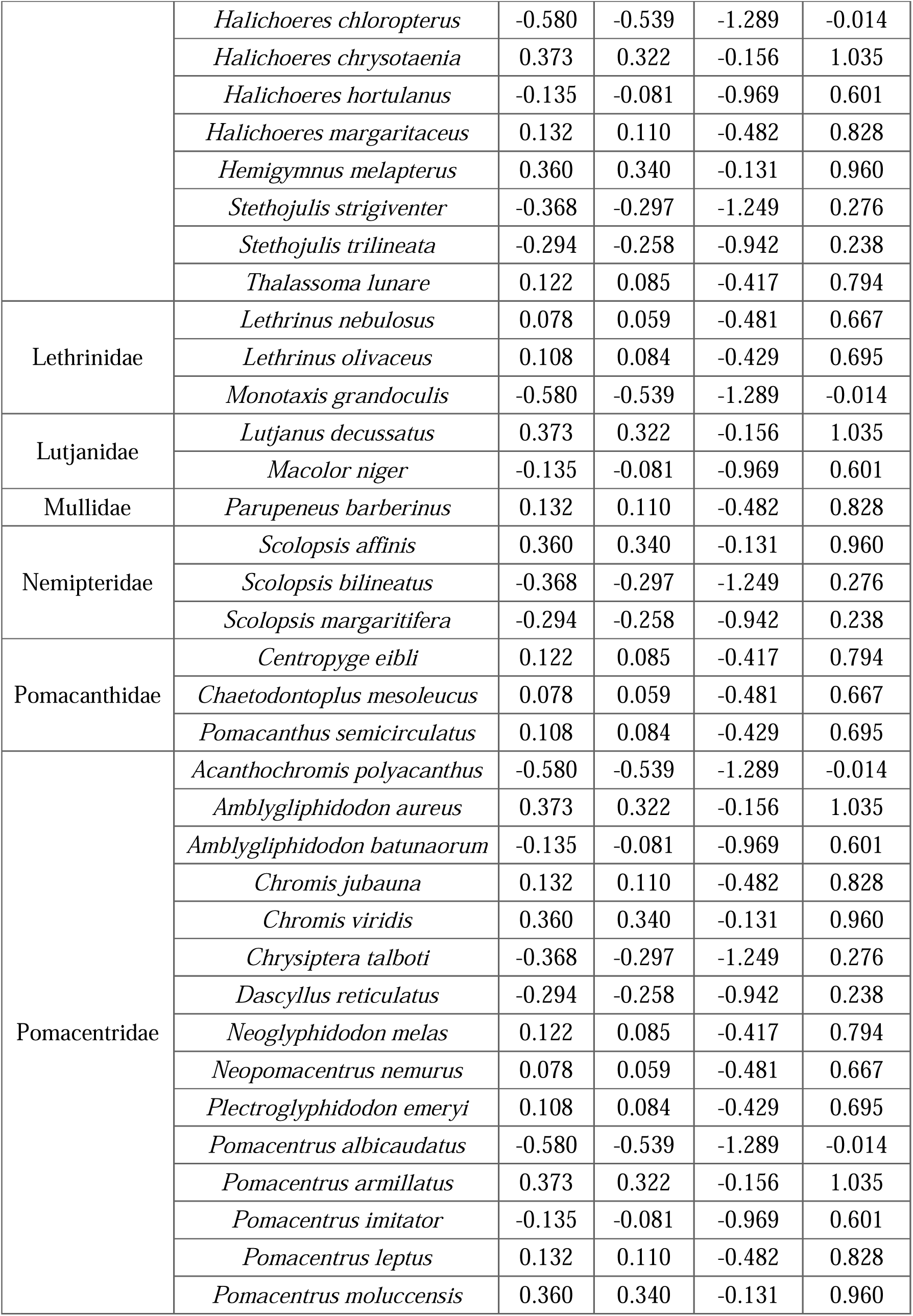
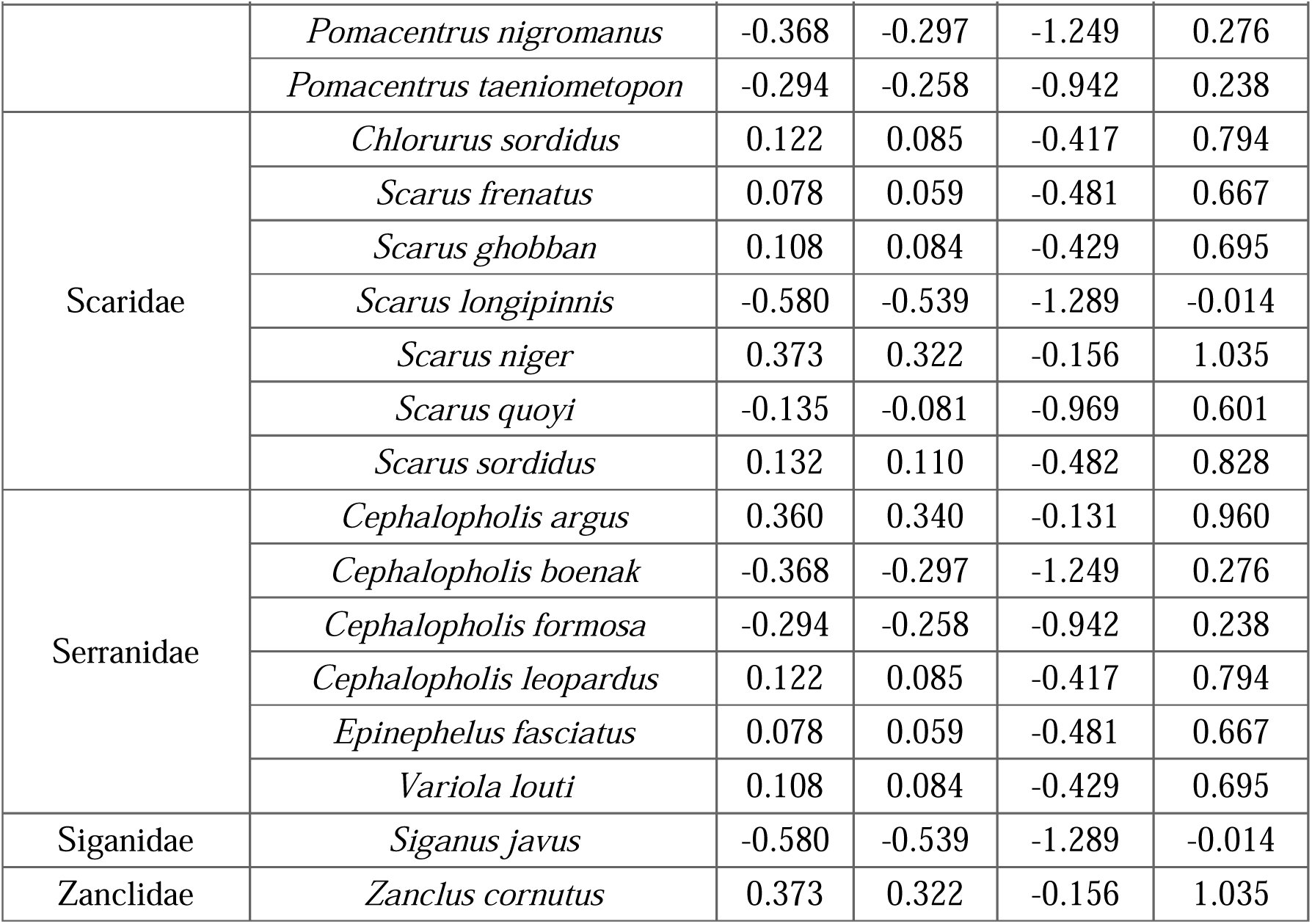
Summary of random intercepts of species for movement time model.

**Table 15:** Summary of species-level random intercepts for precision for movement time model.

| Family | Species | Mean | Median | Lower CI | Upper CI |
| --- | --- | --- | --- | --- | --- |
| Acanthuridae | *Acanthurus albipectoralis* | -1.720 | -1.656 | -3.825 | 0.121 |
|  | *Acanthurus blochii* | -0.019 | -0.074 | -1.915 | 2.202 |
|  | *Acanthurus leucosternon* | 0.064 | 0.043 | -2.508 | 2.618 |
|  | *Acanthurus lineatus* | 0.052 | 0.029 | -2.410 | 2.761 |
|  | *Acanthurus nigrofuscus* | 0.012 | 0.010 | -2.831 | 2.636 |
|  | *Ctenochaetus cyanocheilus* | -0.015 | -0.027 | -2.574 | 2.609 |
|  | *Ctenochaetus strigosus* | -0.043 | -0.034 | -2.624 | 2.560 |
|  | *Naso caesius* | 0.074 | -0.006 | -2.581 | 2.692 |
|  | *Naso elegans* | 0.120 | 0.048 | -2.618 | 2.834 |
|  | *Zebrasoma desjardini* | -0.275 | -0.070 | -3.541 | 2.476 |
| Balistidae | *Balistapus undulatus* | -0.058 | -0.033 | -2.374 | 2.146 |
|  | *Balistoides viridescens* | -0.171 | -0.078 | -3.053 | 2.326 |
|  | *Sufflamen chrysopterum* | 0.002 | -0.018 | -2.430 | 2.731 |
| Caesionidae | *Caesio caerulea* | -0.116 | -0.058 | -3.058 | 2.358 |
|  | *Caesio cuning* | -0.047 | -0.001 | -2.898 | 2.449 |
| Chaetodontidae | *Chaetodon andamanensis* | -1.227 | -1.191 | -2.811 | 0.185 |
|  | *Chaetodon auriga* | 0.037 | 0.044 | -2.434 | 2.481 |
|  | *Chaetodon baronessa* | 0.359 | 0.264 | -1.684 | 2.621 |
|  | *Chaetodon capistratus* | -0.120 | -0.038 | -2.950 | 2.482 |
|  | *Chaetodon decussatus* | -0.073 | -0.085 | -2.549 | 2.371 |
|  | *Chaetodon falcula* | 0.412 | 0.276 | -1.725 | 2.877 |
|  | *Chaetodon lunulatus* | 1.741 | 1.655 | -0.104 | 3.834 |
|  | *Chaetodon meyeri* | 0.091 | 0.003 | -2.393 | 2.946 |
|  | *Chaetodon triangulum* | -0.005 | -0.006 | -2.499 | 2.554 |
|  | *Chaetodon trifacialis* | -0.035 | 0.001 | -2.452 | 2.480 |
| Kyphosidae | *Kyphosus cinerascens* | 0.068 | 0.016 | -2.343 | 2.660 |
| Labridae | *Bodianus mesothorax* | 0.052 | 0.017 | -2.303 | 2.661 |
|  | *Cheilinus abudjubbe* | 0.009 | 0.019 | -2.580 | 2.993 |
|  | *Cheilinus chlorourus* | -0.082 | -0.047 | -2.757 | 2.649 |
|  | *Cheilinus fasciatus* | -0.108 | -0.013 | -3.076 | 2.822 |
|  | *Cheilinus indulatus* | 0.191 | 0.077 | -2.249 | 2.845 |
|  | *Cheilinus trilobatus* | 0.595 | 0.468 | -1.334 | 2.929 |
|  | *Epibulus insidiator* | -0.011 | 0.003 | -2.691 | 2.469 |
|  | *Gomphosus caeruleus* | -0.132 | -0.074 | -2.583 | 2.329 |
|  | *Halichoeres chierchiae* | 0.045 | 0.019 | -2.579 | 2.893 |
|  | *Halichoeres chloropterus* | 0.088 | 0.055 | -2.434 | 2.657 |
|  | *Halichoeres chrysotaenia* | 0.033 | 0.055 | -2.634 | 2.527 |
|  | *Halichoeres hortulanus* | -0.054 | -0.025 | -2.504 | 2.696 |
|  | *Halichoeres margaritaceus* | -0.028 | -0.020 | -2.418 | 2.367 |
|  | *Hemigymnus melapterus* | 0.355 | 0.230 | -1.628 | 2.784 |
|  | *Stethojulis strigiventer* | 0.042 | 0.005 | -2.753 | 2.814 |
|  | *Stethojulis trilineata* | -0.091 | -0.032 | -2.469 | 2.134 |
|  | *Thalassoma lunare* | 0.143 | 0.068 | -2.240 | 2.777 |
| Lethrinidae | *Lethrinus nebulosus* | 0.029 | 0.022 | -2.351 | 2.525 |
|  | *Lethrinus olivaceus* | 0.009 | 0.002 | -2.471 | 2.628 |
|  | *Monotaxis grandoculis* | 0.020 | 0.077 | -2.440 | 2.284 |
| Lutjanidae | *Lutjanus decussatus* | 0.221 | 0.121 | -1.880 | 2.659 |
|  | *Macolor niger* | 0.029 | -0.005 | -2.397 | 2.653 |
| Mullidae | *Parupeneus barberinus* | -0.072 | -0.073 | -2.548 | 2.511 |
| Nemipteridae | *Scolopsis affinis* | 0.114 | 0.036 | -1.901 | 2.577 |
|  | *Scolopsis bilineatus* | 0.684 | 0.590 | -1.358 | 2.897 |
|  | *Scolopsis margaritifera* | -0.136 | -0.089 | -1.970 | 1.626 |
| Pomacanthidae | *Centropyge eibli* | -0.510 | -0.519 | -2.484 | 1.501 |
|  | *Chaetodontoplus mesoleucus* | -0.172 | -0.153 | -2.398 | 1.977 |
|  | *Pomacanthus semicirculatus* | 0.002 | 0.024 | -2.544 | 2.407 |
| Pomacentridae | *Acanthochromis polyacanthus* | 0.079 | 0.083 | -2.426 | 2.411 |
|  | *Amblyglyphidodon aureus* | 0.832 | 0.695 | -1.125 | 3.117 |
|  | *Amblyglyphidodon batunaorum* | -0.140 | -0.027 | -3.105 | 2.282 |
|  | *Chromis jubauna* | 0.052 | -0.001 | -2.976 | 3.154 |
|  | *Chromis viridis* | -0.025 | 0.014 | -2.593 | 2.353 |
|  | *Chrysiptera talboti* | 0.050 | -0.043 | -2.375 | 2.694 |
|  | *Dascyllus reticulatus* | -0.167 | -0.001 | -3.166 | 2.366 |
|  | *Neoglyphidodon melas* | 0.405 | 0.284 | -1.563 | 2.705 |
|  | *Neopomacentrus nemurus* | 0.002 | -0.008 | -2.740 | 2.668 |
|  | *Plectroglyphidodon emeryi* | 0.038 | -0.034 | -2.325 | 2.732 |
|  | *Pomacentrus albicaudatus* | -1.863 | -1.859 | -3.382 | -0.414 |
|  | *Pomacentrus armillatus* | -0.068 | -0.060 | -2.670 | 2.341 |
|  | *Pomacentrus imitator* | 0.497 | 0.390 | -1.613 | 3.189 |
|  | *Pomacentrus leptus* | -0.217 | -0.086 | -3.074 | 2.301 |
|  | *Pomacentrus moluccensis* | 0.009 | -0.029 | -2.641 | 2.751 |
|  | *Pomacentrus nigromanus* | -0.145 | 0.004 | -3.480 | 2.450 |
|  | *Pomacentrus taeniometopon* | 0.019 | 0.022 | -2.440 | 2.535 |
| Scaridae | *Chlorurus sordidus* | 0.272 | 0.224 | -1.235 | 1.977 |
|  | *Scarus frenatus* | -0.004 | 0.015 | -2.988 | 2.905 |
|  | *Scarus ghobban* | -0.041 | -0.003 | -2.713 | 2.669 |
|  | *Scarus longipinnis* | 0.179 | 0.082 | -2.381 | 3.284 |
|  | *Scarus niger* | 0.138 | 0.110 | -2.440 | 2.659 |
|  | *Scarus quoyi* | -0.051 | -0.001 | -2.664 | 2.480 |
|  | *Scarus sordidus* | 0.156 | 0.068 | -2.586 | 3.493 |
| Serranidae | *Cephalopholis argus* | 0.235 | 0.133 | -1.987 | 2.668 |
|  | *Cephalopholis boenak* | 0.004 | -0.004 | -2.498 | 2.506 |
|  | *Cephalopholis formosa* | 0.089 | -0.011 | -2.357 | 2.650 |
|  | *Cephalopholis leopardus* | -0.047 | -0.005 | -2.508 | 2.421 |
|  | *Epinephelus fasciatus* | 0.065 | 0.056 | -2.736 | 2.570 |
|  | *Variola louti* | 0.376 | 0.265 | -1.566 | 2.716 |
| Siganidae | *Siganus javus* | 0.075 | 0.044 | -2.418 | 2.606 |
| Zanclidae | *Zanclus cornutus* | -0.543 | -0.551 | -2.641 | 1.551 |

**Table 16:**
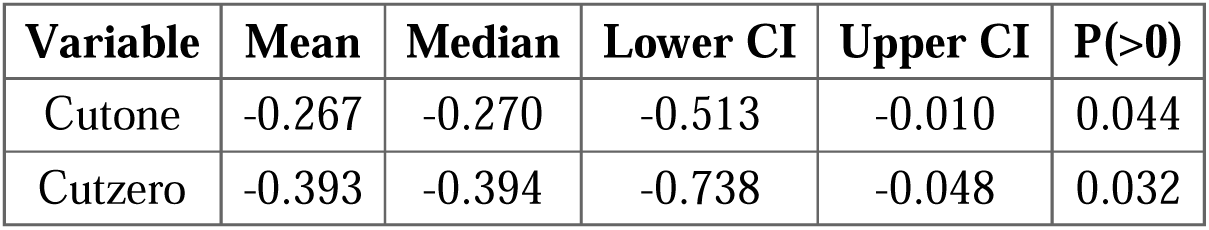
Summary of hyperparameters for movement time mode.

**Table 17:** Summary of fixed effects for movement time model.

| **Variable** | **Mean** | **Median** | **Lower CI** | **Upper CI** | **P(>0)** |
| --- | --- | --- | --- | --- | --- |
| Intercept | -0.168 | -0.153 | -1.050 | 0.674 | 0.370 |
| Protection | -0.082 | -0.075 | -0.688 | 0.543 | 0.425 |
| Treatment (Grouper) | 0.361 | 0.352 | -0.225 | 1.000 | 0.849 |
| Treatment (Negative Control) | -0.185 | -0.176 | -0.816 | 0.437 | 0.313 |
| Treatment (Barracuda) | -0.185 | -0.168 | -0.793 | 0.435 | 0.314 |
| Guild (Invertivore) | 0.056 | 0.062 | -0.649 | 0.708 | 0.574 |
| Rugosity | 0.156 | 0.160 | -0.074 | 0.374 | 0.879 |
| Biomass | -0.054 | -0.050 | -0.286 | 0.168 | 0.339 |
| Group | 0.252 | 0.262 | -0.066 | 0.563 | 0.913 |
| Size class: 10-20 | 0.026 | 0.020 | -0.385 | 0.443 | 0.531 |
| Size class: 20-30 | 0.799 | 0.806 | 0.233 | 1.327 | 0.989 |
| Size class: 30-40 | 0.080 | 0.109 | -0.853 | 0.878 | 0.587 |
| Size class: 40-50 | -0.243 | -0.237 | -0.991 | 0.450 | 0.303 |
| Protection:Treatment (Grouper) | -0.691 | -0.709 | -1.287 | -0.070 | 0.034 |
| Protection:Treatment (Negative Control) | 0.154 | 0.160 | -0.519 | 0.800 | 0.645 |
| Protection:Treatment (Barracuda) | 0.007 | 0.000 | -0.632 | 0.692 | 0.500 |
| Treatment (Grouper):Guild (Invertivore) | 0.153 | 0.145 | -0.510 | 0.807 | 0.649 |
| Treatment (Negative Control):Guild (Invertivore) | -0.240 | -0.242 | -0.872 | 0.413 | 0.274 |
| Treatment (Barracuda):Guild (Invertivore) | -0.425 | -0.428 | -1.055 | 0.193 | 0.131 |
| Protection:Treatment (Negative Control):Guild (Invertivore) | -0.094 | -0.083 | -0.860 | 0.587 | 0.421 |
| Protection:Treatment (Grouper):Guild (Invertivore) | -0.417 | -0.420 | -1.073 | 0.224 | 0.150 |
| Protection:Treatment (Barracuda):Guild (Invertivore) | 0.102 | 0.101 | -0.606 | 0.822 | 0.596 |

**Table 18:** Summary of model fit for movement time model.

| **Type** | **Estimate** | **Est.Error** | **Q2.5** | **Q97.5** |
| --- | --- | --- | --- | --- |
| Conditional | 0.340 | 0.053 | 0.237 | 0.441 |
| Marginal | 0.123 | 0.028 | 0.069 | 0.180 |

## Appendix C: Summary of model for fish bite rates

**Table 1:**
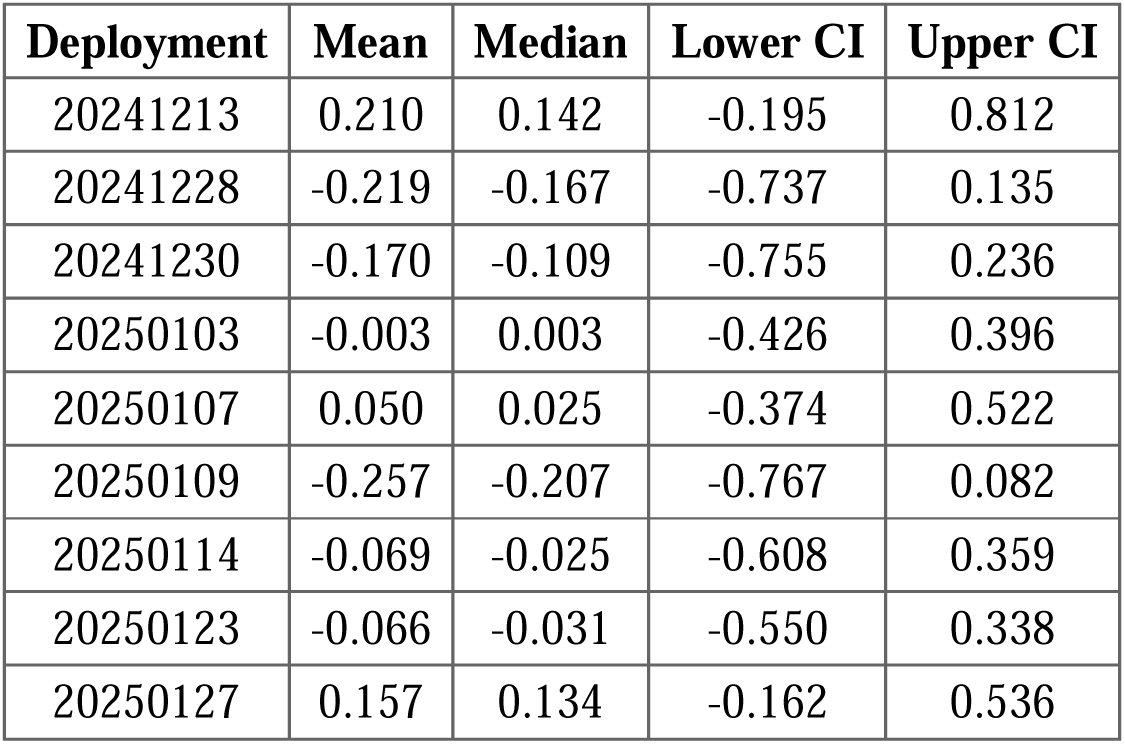
Summary of deployment random effects.

| Deployment | Mean | Median | Lower CI | Upper CI |
| --- | --- | --- | --- | --- |
| 20241213 | 0.210 | 0.142 | -0.195 | 0.812 |
| 20241228 | -0.219 | -0.167 | -0.737 | 0.135 |
| 20241230 | -0.170 | -0.109 | -0.755 | 0.236 |
| 20250103 | -0.003 | 0.003 | -0.426 | 0.396 |
| 20250107 | 0.050 | 0.025 | -0.374 | 0.522 |
| 20250109 | -0.257 | -0.207 | -0.767 | 0.082 |
| 20250114 | -0.069 | -0.025 | -0.608 | 0.359 |
| 20250123 | -0.066 | -0.031 | -0.550 | 0.338 |
| 20250127 | 0.157 | 0.134 | -0.162 | 0.536 |

**Table 2:** Summary of species level random effects.

| Family | Species | Mean | Median | Lower CI | Upper CI |
| --- | --- | --- | --- | --- | --- |
| Acanthuridae | *Acanthurus albipectoralis* | -0.080 | -0.047 | -0.721 | 0.492 |
|  | *Acanthurus blochii* | -0.088 | -0.056 | -0.673 | 0.453 |
|  | *Acanthurus leucosternon* | -0.227 | -0.141 | -1.082 | 0.388 |
|  | *Acanthurus lineatus* | 0.092 | 0.054 | -0.408 | 0.672 |
|  | *Acanthurus nigrofusus* | 0.154 | 0.101 | -0.402 | 0.838 |
| Balistidae | *Sufflamen chrysopteron* | 0.032 | 0.009 | -0.505 | 0.603 |
| Chaetodontidae | *Chaetodon andamanensis* | -0.090 | -0.067 | -0.619 | 0.419 |
|  | *Chaetodon baronessa* | -0.048 | -0.029 | -0.585 | 0.473 |
|  | *Chaetodon falcula* | -0.068 | -0.043 | -0.607 | 0.438 |
|  | *Chaetodon lunulatus* | 0.386 | 0.357 | -0.045 | 0.951 |
|  | *Chaetodon trifacialis* | -0.060 | -0.033 | -0.622 | 0.473 |
| Labridae | *Cheilinus abudjubbe* | 0.129 | 0.077 | -0.355 | 0.723 |
|  | *Cheilinus trilobatus* | -0.148 | -0.087 | -0.907 | 0.459 |
|  | *Halichoeres chierchiae* | -0.074 | -0.044 | -0.736 | 0.514 |
|  | *Halichoeres chrysotaenia* | 0.185 | 0.118 | -0.318 | 0.866 |
|  | *Halichoeres hortulanus* | -0.236 | -0.170 | -0.960 | 0.299 |
|  | *Hemigymnus melapterus* | -0.140 | -0.084 | -0.872 | 0.464 |
|  | *Stethojulis strigiventer* | -0.152 | -0.090 | -0.861 | 0.421 |
|  | *Thalassoma lunare* | -0.053 | -0.027 | -0.703 | 0.535 |
| Mullidae | *Parupeneus barberinus* | -0.143 | -0.081 | -0.852 | 0.421 |
| Pomacanthidae | *Centropyge eibli* | 0.009 | 0.006 | -0.434 | 0.456 |

| **Family** | **Species** | **Mean** | **Median** | **Lower CI** | **Upper CI** |
| --- | --- | --- | --- | --- | --- |
|  | *Chaetodontoplus mesoleucus* | -0.147 | -0.088 | -0.874 | 0.417 |
|  | *Pomacanthus semicirculatus* | -0.180 | -0.111 | -0.939 | 0.396 |
| Pomacentridae | *Chromis jubauna* | 0.067 | 0.024 | -0.477 | 0.737 |
|  | *Pomacentrus albicaudatus* | 0.229 | 0.194 | -0.177 | 0.746 |
|  | *Pomacentrus leptus* | -0.156 | -0.090 | -0.875 | 0.421 |
|  | *Pomacentrus nigromanus* | 0.345 | 0.263 | -0.153 | 1.102 |
| Scaridae | *Chlorurus sordidus* | -0.283 | -0.235 | -0.921 | 0.166 |
|  | *Scarus frenatus* | 0.290 | 0.233 | -0.171 | 0.945 |
|  | *Scarus ghobban* | 0.166 | 0.099 | -0.372 | 0.855 |
|  | *Scarus niger* | 0.058 | 0.037 | -0.470 | 0.608 |
|  | *Scarus quoyi* | -0.002 | -0.003 | -0.528 | 0.539 |
|  | *Scarus sordidus* | 0.204 | 0.129 | -0.330 | 0.952 |
| Serranidae | *Cephalopholis argus* | -0.186 | -0.114 | -0.979 | 0.397 |
| Zanclidae | *Zanclus cornutus* | -0.237 | -0.161 | -1.006 | 0.315 |

**Table 3:**
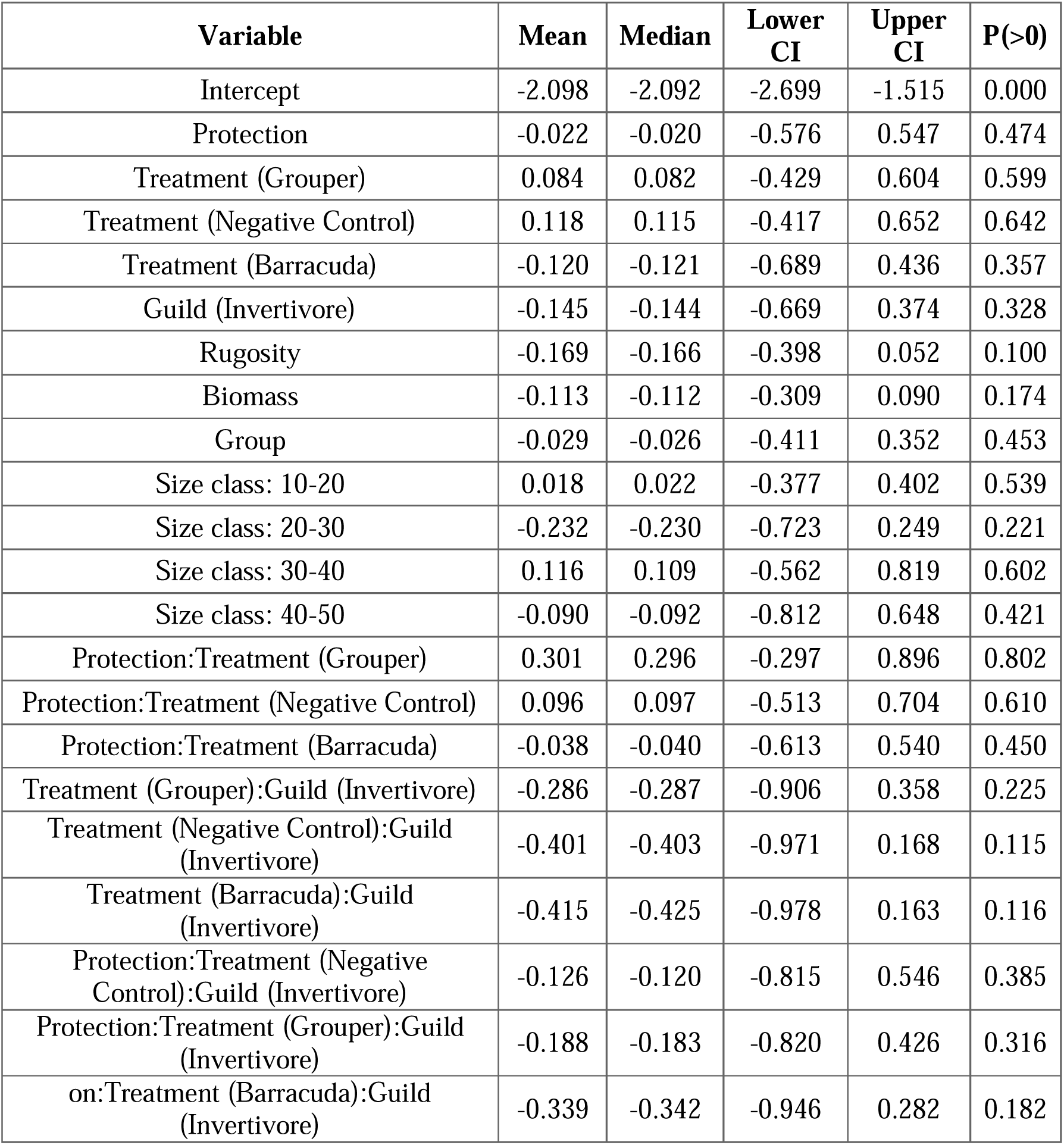
Summary of fixed effects.

**Table 4:** Summary of model fit.

| Type | Estimate | Est.Error | Q2.5 | Q97.5 |
| --- | --- | --- | --- | --- |
| Conditional | 0.316 | 0.090 | 0.156 | 0.503 |
| Marginal | 0.216 | 0.083 | 0.084 | 0.400 |

## Appendix D: Summary of effect of predator decoy treatments and controls on fish behavioural responses

**Table 1:** Comparison of positive and negative control effects (log odds ratios) on fish time budgets.

| **Behaviour** | **Median** | **90% Lower CI** | **50% Lower CI** | **50% Upper CI** | **90% Upper CI** | **P(>0)** |
| --- | --- | --- | --- | --- | --- | --- |
| Foraging | 0.052 | -1.327 | -0.535 | 0.679 | 1.576 | 0.525 |
| Movement | 0.383 | -1.178 | -0.191 | 0.944 | 1.747 | 0.675 |
| Vigilance | -0.994 | -2.396 | -1.553 | -0.419 | 0.545 | 0.133 |

**Table 2:**
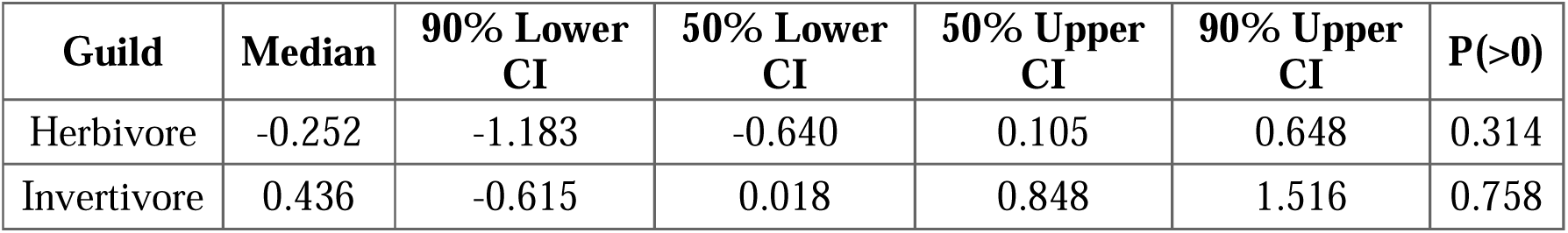
Comparison of positive and negative control effects (log response ratios) on the bite rates of fish.

**Table 3:**
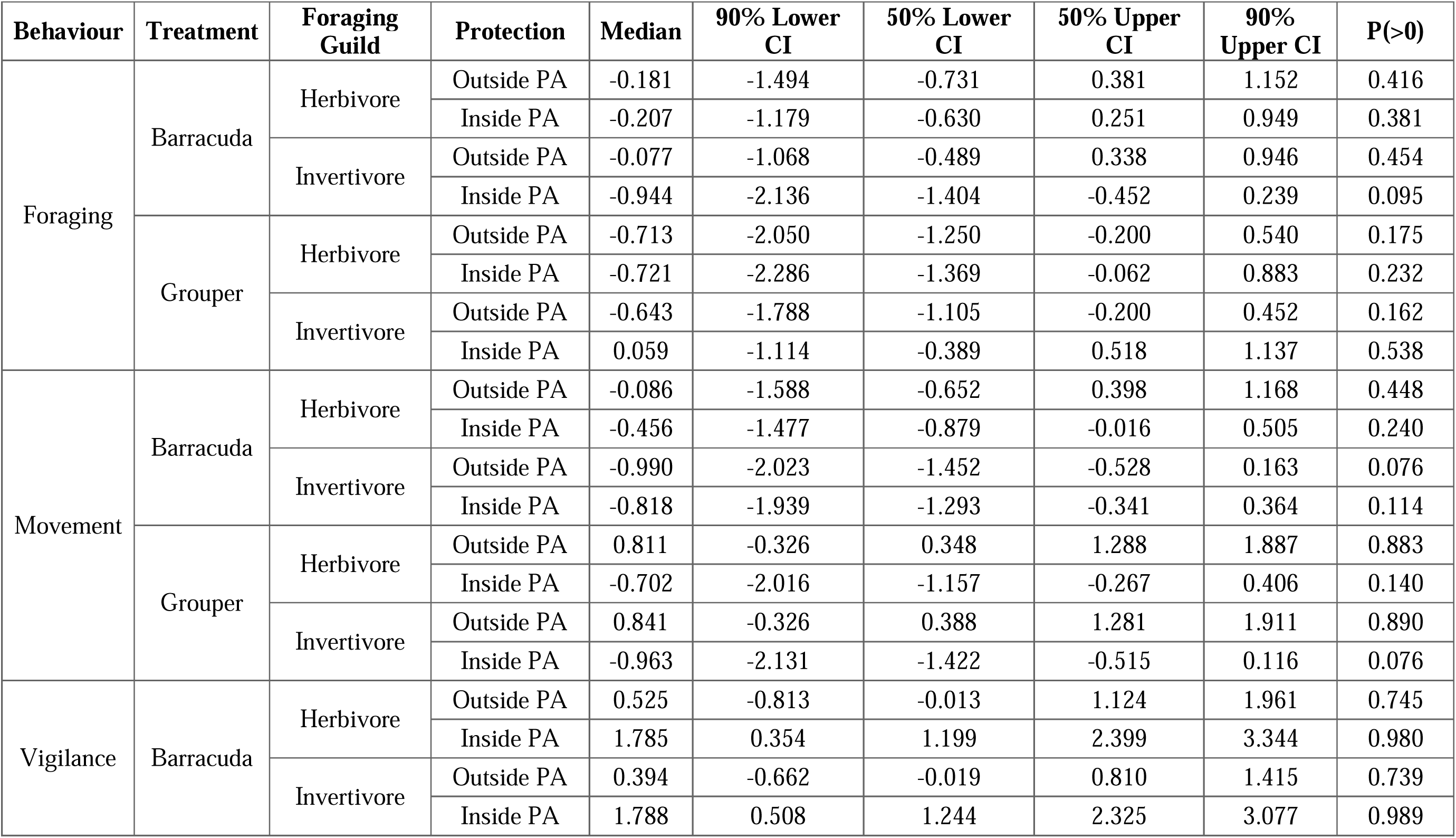
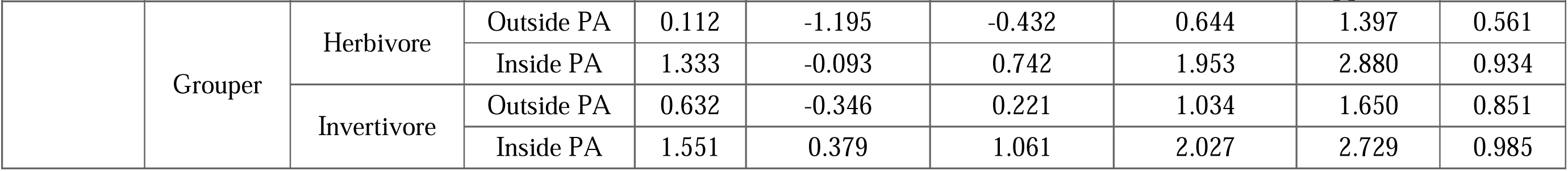
Comparison of treatment effects (log odds ratios) on fish time budgets within and outside protected areas across guilds.

| Behaviour | Treatment | Foraging Guild | Protection | Median | 90% Lower CI | 50% Lower CI | 50% Upper CI | 90% Upper CI | P(>0) |
| --- | --- | --- | --- | --- | --- | --- | --- | --- | --- |
| Foraging | Barracuda | Herbivore | Outside PA | -0.181 | -1.494 | -0.731 | 0.381 | 1.152 | 0.416 |
|  |  |  | Inside PA | -0.207 | -1.179 | -0.630 | 0.251 | 0.949 | 0.381 |
|  |  | Invertivore | Outside PA | -0.077 | -1.068 | -0.489 | 0.338 | 0.946 | 0.454 |
|  |  |  | Inside PA | -0.944 | -2.136 | -1.404 | -0.452 | 0.239 | 0.095 |
|  | Grouper | Herbivore | Outside PA | -0.713 | -2.050 | -1.250 | -0.200 | 0.540 | 0.175 |
|  |  |  | Inside PA | -0.721 | -2.286 | -1.369 | -0.062 | 0.883 | 0.232 |
|  |  | Invertivore | Outside PA | -0.643 | -1.788 | -1.105 | -0.200 | 0.452 | 0.162 |
|  |  |  | Inside PA | 0.059 | -1.114 | -0.389 | 0.518 | 1.137 | 0.538 |
| Movement | Barracuda | Herbivore | Outside PA | -0.086 | -1.588 | -0.652 | 0.398 | 1.168 | 0.448 |
|  |  |  | Inside PA | -0.456 | -1.477 | -0.879 | -0.016 | 0.505 | 0.240 |
|  |  | Invertivore | Outside PA | -0.990 | -2.023 | -1.452 | -0.528 | 0.163 | 0.076 |
|  |  |  | Inside PA | -0.818 | -1.939 | -1.293 | -0.341 | 0.364 | 0.114 |
|  | Grouper | Herbivore | Outside PA | 0.811 | -0.326 | 0.348 | 1.288 | 1.887 | 0.883 |
|  |  |  | Inside PA | -0.702 | -2.016 | -1.157 | -0.267 | 0.406 | 0.140 |
|  |  | Invertivore | Outside PA | 0.841 | -0.326 | 0.388 | 1.281 | 1.911 | 0.890 |
|  |  |  | Inside PA | -0.963 | -2.131 | -1.422 | -0.515 | 0.116 | 0.076 |
| Vigilance | Barracuda | Herbivore | Outside PA | 0.525 | -0.813 | -0.013 | 1.124 | 1.961 | 0.745 |
|  |  |  | Inside PA | 1.785 | 0.354 | 1.199 | 2.399 | 3.344 | 0.980 |
|  |  | Invertivore | Outside PA | 0.394 | -0.662 | -0.019 | 0.810 | 1.415 | 0.739 |
|  |  |  | Inside PA | 1.788 | 0.508 | 1.244 | 2.325 | 3.077 | 0.989 |
|  | Grouper | Herbivore | Outside PA | 0.112 | -1.195 | -0.432 | 0.644 | 1.397 | 0.561 |
|  |  |  | Inside PA | 1.333 | -0.093 | 0.742 | 1.953 | 2.880 | 0.934 |
|  |  | Invertivore | Outside PA | 0.632 | -0.346 | 0.221 | 1.034 | 1.650 | 0.851 |
|  |  |  | Inside PA | 1.551 | 0.379 | 1.061 | 2.027 | 2.729 | 0.985 |

**Table 4:** Comparison of treatment effects (log response ratios) on the bite rates of fish within and outside protected areas across guilds.

| Foraging Guild | Treatment | Protection | Median | 90% Lower CI | 50% Lower CI | 50% Upper CI | 90% Upper CI | P(>0) |
| --- | --- | --- | --- | --- | --- | --- | --- | --- |
| Herbivore | Barracuda | Outside PA | -0.175 | -0.994 | -0.492 | 0.159 | 0.629 | 0.357 |
|  |  | Inside PA | -0.227 | -1.044 | -0.555 | 0.109 | 0.583 | 0.323 |
|  | Grouper | Outside PA | 0.118 | -0.619 | -0.191 | 0.434 | 0.871 | 0.599 |
|  |  | Inside PA | 0.562 | -0.328 | 0.170 | 0.938 | 1.471 | 0.834 |
| Invertivore | Barracuda | Outside PA | -0.761 | -1.585 | -1.112 | -0.438 | 0.031 | 0.055 |
|  |  | Inside PA | -1.322 | -2.142 | -1.667 | -0.971 | -0.437 | 0.008 |
|  | Grouper | Outside PA | -0.289 | -1.256 | -0.700 | 0.107 | 0.703 | 0.312 |
|  |  | Inside PA | -0.131 | -0.849 | -0.417 | 0.158 | 0.582 | 0.376 |

